# Separating neural oscillations from aperiodic 1/f activity: challenges and recommendations

**DOI:** 10.1101/2021.10.15.464483

**Authors:** Moritz Gerster, Gunnar Waterstraat, Vladimir Litvak, Klaus Lehnertz, Alfons Schnitzler, Esther Florin, Gabriel Curio, Vadim Nikulin

## Abstract

Electrophysiological power spectra typically consist of two components: An aperiodic part usually following an 1/f power law *P*∝1/*f*^β^and periodic components appearing as spectral peaks. While the investigation of the periodic parts, commonly referred to as neural oscillations, has received considerable attention, the study of the aperiodic part has only recently gained more interest. The periodic part is usually quantified by center frequencies, powers, and bandwidths, while the aperiodic part is parameterized by the y-intercept and the 1/f exponent β. For investigation of either part, however, it is essential to separate the two components.

In this article, we scrutinize two frequently used methods, FOOOF (Fitting Oscillations & One-Over-F) and IRASA (Irregular Resampling Auto-Spectral Analysis), that are commonly used to separate the periodic from the aperiodic component. We evaluate these methods using diverse spectra obtained with electroencephalography (EEG), magnetoencephalography (MEG), and local field potential (LFP) recordings relating to three independent research datasets. Each method and each dataset poses distinct challenges for the extraction of both spectral parts. The specific spectral features hindering the periodic and aperiodic separation are highlighted by simulations of power spectra emphasizing these features. Through comparison with the simulation parameters defined a priori, the parameterization error of each method is quantified. Based on the real and simulated power spectra, we evaluate the advantages of both methods, discuss common challenges, note which spectral features impede the separation, assess the computational costs, and propose recommendations on how to use them.

## Introduction

Analysis of macroscopic electromagnetic brain activity (e.g., by EEG and MEG) has long been focusing on the investigation of ‘rhythmic’ neural oscillations. In the frequency domain, neural oscillations appear as distinct spectral peaks, also referred to as the periodic part of the spectrum (Buzsáki & Draguhn, 2004; Engel et al., 2001; Schnitzler & Gross, 2005). The full spectrum, however, also consists of a continuous component whose analysis has, so far, seen less attention. This aperiodic or ‘arrhythmic’ part of the spectrum (Freeman & Zhai, 2009; Miller et al., 2009) has been related to the integration of underlying synaptic currents (Buzsáki et al., 2012). Since the time series of the aperiodic part is typically self-similar across many temporal scales, it is also referred to as “fractal” or “scale-free” activity. The power spectral density (PSD) of the aperiodic component follows a power law *P*∝1/*f*^β^(Miller et al., 2009) and is sometimes called 1/f activity for that reason. In this text, we will refer to the scaling exponent β in this equation as 1/f exponent.

The investigation of neural oscillations has received much attention in electrophysiological studies (Buzsáki & Draguhn, 2004; Singer, 1999; Ward, 2003). However, the standard analysis of assessing periodic power through bandpass filtering is problematic because the pass-band comprises both periodic and aperiodic activity. If the power of aperiodic activity changes between two conditions, analyzing neural oscillations in bandpass filtered signals would hence be confounded by these changes in the aperiodic part of the spectra. For that reason, estimating the 1/f component before determining the power of periodic activity has recently been suggested (Donoghue et al., 2021; Wen & Liu, 2016).

Besides investigating neural oscillations, the investigation of the aperiodic component has recently gained considerable interest (He, 2014; Kello et al., 2010). For example, the 1/f exponent was shown to change with task (Waschke et al., 2021), age (He et al., 2019; Schaworonkow & Voytek, 2021; Voytek et al., 2015), and disease (Molina et al., 2020; Robertson et al., 2019) and it decreases with cortical depth (Halgren et al., 2021). Furthermore, using computational modeling, (Gao et al., 2017) suggested the 1/f exponent β as an estimator of excitation–inhibition (E–I) balance. Many studies comparing conscious states—associated with increased excitation—to unconscious states, such as NREM sleep (Lendner et al., 2020; Miskovic et al., 2019) and anesthesia (Colombo et al., 2019; Waschke et al., 2021)—typically associated with pronounced inhibitory processes—seem to support this concept.

But how to best estimate the 1/f exponent? This will be the main question discussed in this study. One option is to simply fit a straight line using (robust) linear regression. (Gao et al., 2017) used this method in the frequency ranges apart from pronounced oscillatory peaks in electrocorticography (ECoG) data and identified distinct 1/f exponents during wakefulness and anesthesia. However, in the presence of periodic components, this method is error-prone because larger periodic peaks will bias the linear regression fit.

Irregular-resampling autospectral analysis (IRASA) (Wen & Liu, 2016) aims to separate periodic components from the aperiodic part of the spectrum. Due to their fractal nature, aperiodic time series remain robust against resampling, whereas periodic components are strongly affected by this procedure. IRASA takes advantage of this dichotomy and ‘removes’ the periodic parts from a spectrum. The—ideally—pure aperiodic part of the spectrum obtained with this method can then be used for fitting the 1/f exponent.

Another method, ‘fitting oscillations & one over f’ (FOOOF) (Donoghue et al., 2020), aims at modeling the periodic components: It iteratively applies Gaussian fits to all periodic components and hereby obtains a model of the periodic part. This model of periodic activity is subtracted from the spectrum to obtain an—ideally—pure aperiodic component which can be used for fitting β. In addition, the periodic model allows for analyzing the periodic components (e.g., regarding center frequencies, bandwidths, and power) without the bias from aperiodic activity.

This article highlights and discusses the general challenges of estimating 1/f exponents. In addition, we also discuss method-specific challenges of FOOOF and IRASA, the most commonly used algorithms for that purpose.

In the <u>Methods</u> section, we will introduce our simulations, our datasets, and both algorithms FOOOF and IRASA. We will analyze challenges by the example of FOOOF in section <u>FOOOF</u> and by the example of IRASA in the section <u>IRASA</u>. To aim for broad applicability of our assessment, we will apply these methods to simulations with known ground truth in addition to various electrophysiological signals obtained from empirical EEG, gradiometer MEG, magnetometer MEG, source-reconstructed voxel activity from MEG, and subthalamic nucleus-(STN-)LFP data acquired by three independent research groups. We will discuss these challenges in the section <u>Discussion</u>, and we will provide some guidance on how to use these methods in the <u>Conclusion</u> section.

## Methods

### Simulations

We simulate aperiodic 1/f activity by constructing a Fourier power spectrum following a preset 1/*f*^β^power-law. The corresponding phases of the Fourier spectrum are distributed uniformly randomly. To add oscillations, we add Gaussian-shaped peaks to the Fourier power spectrum with amplitudes *A* and a spectral extent given by center frequencies *f*_*center*_ and variances 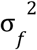. The corresponding time series, consisting of both ‘neural’ oscillations and aperiodic activity, is then obtained by applying the inverse fast Fourier transform. The simulated time series either have a duration of 180 s at a sampling rate of *f*_*sample*_ *=* 2400 Hz or are matched to the empirical data to which a simulation might be compared. If noted in the text, Gaussian white noise may be added to the time series afterward. Since most algorithms to generate 1/f activity lead to identical power spectra, the specific choice of the algorithm has no impact on the present analysis.

### Empirical data

We compare the results from our simulations to three empirical datasets.

#### Dataset 1

Dataset 1 was re-analyzed from (Litvak et al., 2010, 2011) and contains MEG and LFP data of 14 Parkinsonian patients after bilateral implantation of subthalamic nucleus (STN) stimulation electrodes (Medtronic, Minneapolis, MN, USA with four platinum-iridium cylindrical surfaces of diameter 1.27 mm, length 1.5 mm, and center-to-center separation 2 mm) for deep brain stimulation (DBS). The joint ethics committee of the National Hospital of Neurology and Neurosurgery and the University College London Institute of Neurology approved the study, and all patients gave their written informed consent. The patients were recorded three days after surgery when the electrode leads were still externalized. The recordings were obtained during a Parkinsonian state OFF medication (after overnight withdrawal) and an ON medication state. MEG (275 channels, CTF/VSM MedTech, Vancouver, Canada) and DBS-LFP were recorded simultaneously during three minutes of resting-state at a sampling rate of *f*_*sample*_ *=* 2400 Hz. The LFP recordings were referenced to the right mastoid during recording and later re-referenced to a bipolar montage between adjacent electrode contacts. This results in 3 bipolar LFP channels per hemisphere. All data were bandpass filtered in hardware between 1–600 Hz. MEG source reconstruction was performed with varying regularization by Linearly Constrained Minimum Variance beamformer (Van Veen et al., 1997). Aside from the six LFP channels, the dataset contains three MEG channels per patient from voxels located in the supplementary motor area (SMA), left primary motor cortex (M1), and right M1. In this study, we draw examples from voxel data located in the supplementary motor area (SMA) of patients 5 and 6 and bipolarly recorded LFPs from the STN of patients 9 and 10. Details regarding the data recording, processing, and inverse modeling can be obtained from the original publications of this dataset (Litvak et al., 2010, 2011). In this study, we further process this dataset by applying a notch filter at 100, 150, …, 600 Hz power line noise for visualization purposes (multi-taper estimation of sinusoidal components “spectrum_fit” of MNE python (Gramfort et al., 2013)). Note, that a notch filter should not be applied to frequency ranges used for FOOOF fitting. Therefore, we exclude 50 Hz from the notch filter (see SI Fig. 1)).

#### Dataset 2

Dataset 2 contains EEG data from a 12-year-old boy with absence epilepsy recorded at the Department of Epileptology at the University of Bonn (Gerster et al., 2020). The university’s ethics committee approved the study, and a parent gave written informed consent that the clinical data might be used and published for research purposes. EEG data were acquired at a sampling rate of *f*_*sample*_ *=* 256 Hz (16-bit A/D conversion) within a bandwidth of 0.3–70 Hz from 19 electrodes in a bipolar montage. The locations and nomenclature of these electrodes are standardized by the American Electroencephalographic Society (Sharbrough, 1991). The EEG was recorded over several hours and contains 5 absence seizures. In this study, we present 40 s of the bipolar EEG channel “F3−C3” during one absence seizure.

#### Dataset 3

Dataset 3 contains MEG recorded with gradiometers and magnetometers and LFP data from a Parkinsonian patient recorded at the Universitätsklinikum Düsseldorf. The data were acquired using a whole-head MEG system with 306 channels (Elekta Vectorview, Elekta Neuromag, Finland), and segmented “1–3–3–1” electrode DBS-LFP (Abbott St. Jude Medical model 6172, contact height: 1.5 mm with 0.5 mm vertical spacing) during the ON- and OFF-medication state (after overnight withdrawal). The patient was recorded 1 day after surgery when the electrode leads were still externalized. The resting-state was recorded for 10 min at a sample rate of *f*_*sample*_ *=* 2400 Hz. The LFP recordings were referenced to the right mastoid during recording and later re-referenced to a bipolar montage between adjacent electrodes. The data were offline band-pass filtered between 0.3 Hz and 600 Hz and notch-filtered at 50, 100, …, 600 Hz power line noise (with a second-order IIR filter of bandwidth 1 Hz). Note that notch filtering in the fitting range at 50 Hz is unproblematic with using IRASA. The patient gave written consent to participate in the study, which was approved by the Ethics committee of the Universitätsklinikum Düsseldorf. In this study, we analyze data from one gradiometer channel, one magnetometer channel, and one LFP channel of the subject.

### Power spectral densities (PSDs)

We calculate the PSDs from the simulated and recorded time series using the Welch algorithm. We use a segment length of 1 s which corresponds to a frequency resolution of 1 Hz, and the Hann-windowed segments overlap by 50%. Please note that other segment lengths can be used depending on the properties of the data. However, for FOOOF, the PSDs should be sufficiently smooth to avoid fitting noise peaks. IRASA receives time series as input and calculates the PSDs internally. For IRASA, the PSD resolution should be sufficiently high. We, therefore, use a segment length of 4 seconds (corresponding to a resolution of 0.25 Hz), Hann windows, and 50% overlap.

### Irregular-Resampling Auto-Spectral Analysis (IRASA)

Irregular-resampling auto-spectral analysis (IRASA) aims at separating periodic components from the aperiodic part of the signal (Wen & Liu, 2016). In contrast to FOOOF, the algorithm requires time series as input (Fig. 1 a) and does not explicitly model the signals’ spectra. The input time series is upsampled by a set of predefined resampling factors *h*_*i*_ ∈ *h*_*set*_. By default, *h*_*set*_ ranges from 1.1 to 1.9 with increments of 0.05, yielding 17 resampling factors *h*_*set*_ *=* {1. 1, 1. 15, …, 1. 9}. In addition, the time series is downsampled by all inverse resampling factors 1/*h*_*i*_, with *h*_*i*_ ∈ *h*_*set*_. For each of the 17 pairs of up- und downsampled spectra (Fig. 1 b), the geometric mean of the PSD is calculated (Fig. 1 c). For illustration purposes in Fig. 1, we use a very small *h*_*set*_ *=* {1. 3, 1. 6, 2}. Finally, the median is calculated from all 17 geometric means, yielding the aperiodic component (Fig. 1 d). The compound oscillatory part of the spectrum is obtained by subtracting the aperiodic component from the original PSD. After applying IRASA, the slope β can be obtained by fitting the aperiodic component in double logarithmic space in the predefined fitting range.

**Fig. 1.**
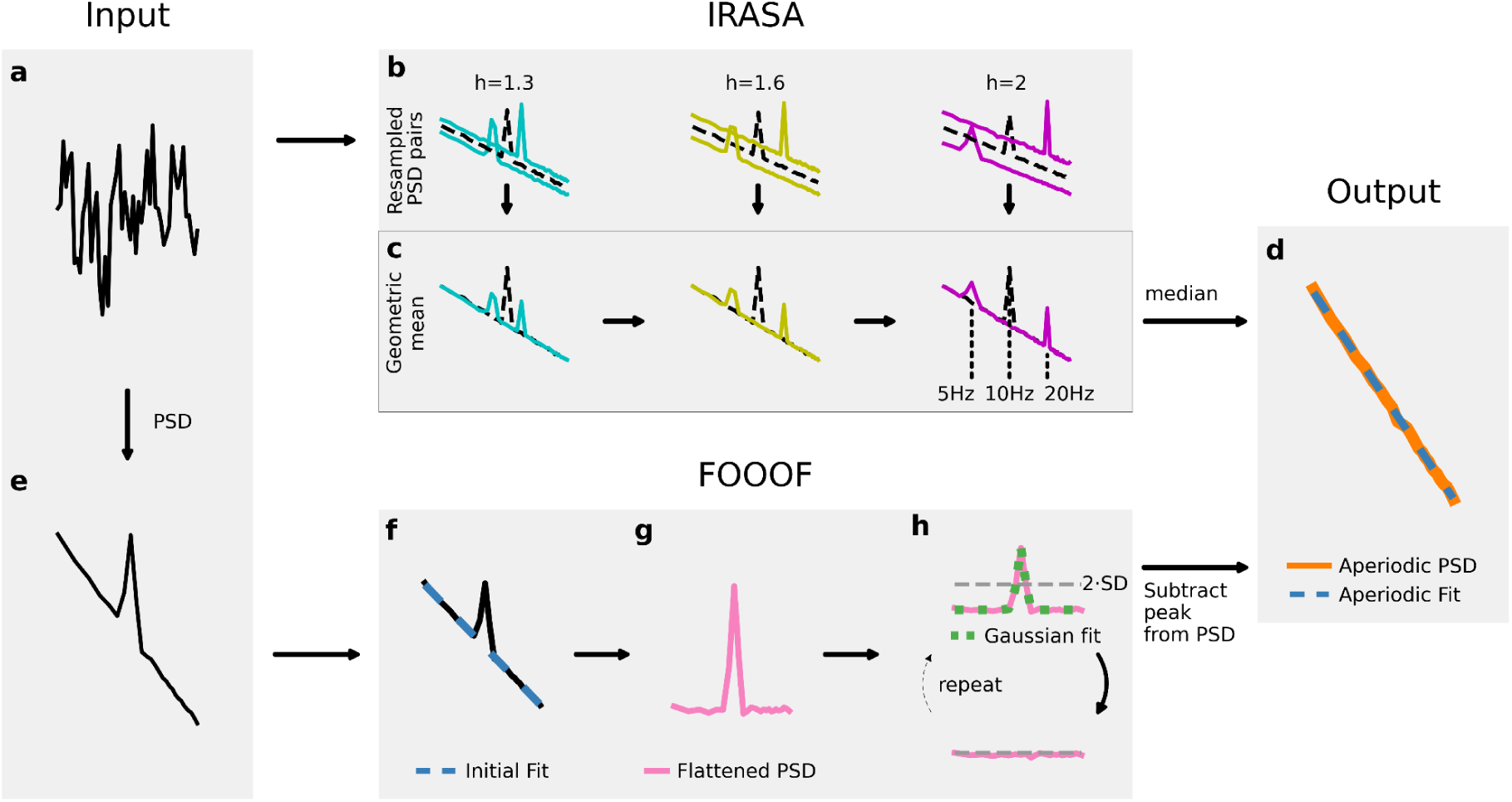
*Algorithms for 1/f estimation*. IRASA: *a) Simulated time series. b) PSDs of resampled time series on the y-axis and frequencies on the x-axis. In this figure, the time series is upsampled by the resampling factors h*_*i*_ *of the h*_*set*_ *=* {1. 3, 1. 6, 2} *and downsampled by* 1/*h*_*i*_. *c) The geometric mean of all resampling pairs (h*_*i*_, 1/*h*_*i*_*) is calculated. d) The aperiodic component (orange) is the median of the geometric means. A final fit (dashed-blue) estimates the y-intercept and the 1/f exponent* β. FOOOF: *e) A PSD is calculated from the time series. f) FOOOF applies an initial linear fit (dashed-blue) to the PSD in log-log space and g) subtracts the obtained linear trend from the spectrum. h) A Gaussian model (dotted-green) is fitted to the largest peak exceeding the thresholds (dashed-grey) and removes it. The relative threshold is recalculated from the peak-removed flattened spectrum (pink). The procedure is repeated until no peak exceeds the relative threshold. d) Subtraction of all Gaussian models from the original PSD yields the aperiodic component, which is then finally re-fit*

As parameters, IRASA requires the fitting range, the resampling factors *h*_*set*_, and the segment length for the PSD calculation. In this study, we vary the fitting range and the *h*_*set*_ but keep the segment length at 4 s. IRASA’s Python implementation used for this article was adapted from the YASA toolbox (Vallat, 2019) and is published along with the complete code for this study on GitHub at https://github.com/moritz-gerster/oscillation_and_1-f_separation.

### Fitting-oscillations-&-one-over-f (FOOOF)

FOOOF was introduced to parameterize neural power spectra as a combination of an aperiodic component and peaks representing oscillatory processes (Donoghue et al., 2020). The Python-based toolbox works as outlined in Fig. 1. First, the PSD of the time-series of interest (Fig. 1 a) is calculated and input into the algorithm, Fig. 1 e). Next, FOOOF calculates an initial robust linear fit of the spectrum in double logarithmic space, Fig. 1 f), and subtracts the result from the spectrum, Fig. 1 g). In this flattened spectrum, a relative threshold is calculated based on the standard deviation (SD) of the spectrum, Fig. 1 h). The relative threshold is set to two times the SD, by default *thresh*_*rel*_ *=* 2 *SD*. Optionally, FOOOF also allows setting an additional absolute threshold for the peak heights, but it is set to 0 by default (*thresh*_*abs*_ *=* 0). A Gaussian function is fitted to the largest peak of the flattened PSD exceeding both thresholds and then subtracted from the spectrum. Note, that this fit is not applied to negative peaks in the spectrum subceeding both thresholds. Therefore, spectral dips caused by notch filtering should be avoided. This procedure is iterated for the next largest peak after subtracting the previous peak until no peaks are exceeding the thresholds. The oscillatory components are finally obtained by fitting a multivariate Gaussian to all extracted peaks simultaneously. After the iterations, the initial fit is added back to the flattened peak-free PSD, which results in the aperiodic component of the PSD, Fig. 1 d). Afterward, this aperiodic component is fitted again, leading to the final fit with y-intercept and slope β as parameters. The fitted Gaussian functions are parameterized by center frequency *f*_*center*_(“CF”), amplitude *A*_*f*_ (“PW”), and bandwidth 2 · 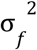 (“BW”).

The algorithm can be used with the following parameters: “peak_width_limits” allows setting the minimum and maximum peak width limits of the Gaussian fits. The default is set to 0.5 Hz and 12 Hz, respectively. “max_n_peaks’’ determines the maximum number of peak fitting iterations. The default is set to infinity. “min_peak_height” is the absolute threshold and corresponds to the smallest peaks that will be fitted in the units of the input data. The absolute threshold is set to 0 by default. “peak_threshold” is the relative threshold in SD multiples that peaks must exceed to be fitted and defaults to “peak_threshold” = 2 SD. The peak fitting stops when all remaining peaks are below either of these two thresholds. “aperiodic mode” allows for two modes of modeling: “fixed” and “knee” which allows modeling a bend (i.e., a “knee”) in the PSD of the aperiodic component. The default is “fixed.” Finally, FOOOF accepts a fitting range for which the algorithm performs the given steps. For a detailed description of this algorithm, we refer the reader to the methods section of the original publication (Donoghue et al., 2020).

In this study, we keep FOOOF at the default parameters if not stated otherwise and input PSDs with a spectral resolution of 1Hz.

## FOOOF

### Challenge 1: The spectral plateau disrupts the 1/f power law

The 1/f power law is sometimes called “scale-free” because log-transformed power is typically approaching a linear function across an extended range of log-transformed frequencies. For electrophysiological PSDs, however, this concept should be exercised carefully. For example, (He et al., 2010) measured different values for the slope β for frequency ranges 0.01–0.1 Hz and 1–100 Hz and found a small plateau in the range from 0.1 Hz to 1 Hz. In many studies, these low-frequency ranges <0.1 Hz are eliminated by a hardware high-pass filter. However, this finding underlines the importance of selecting a representative frequency range to fit the 1/f slope.

In addition to the aforementioned low-frequency plateau, one regularly encounters a high-frequency spectral plateau (or flattening) in spectra of electrophysiological data. Such plateaus might be due to the presence of Gaussian noise which appears as a horizontal line with a slope β *=* 0 in double-logarithmic space and disrupts the 1/f power law. The origin of such white noise is often due to EMG artifacts and electronic noise of the recording system (Waterstraat et al., 2015). It has been shown in EEG (Scheer et al., 2006; Waterstraat, Burghoff, et al., 2015) and MEG (Waterstraat et al., 2021) that extremely low-noise recording devices can shift this high-frequency plateau into the kHz range—leaving a wider unaffected frequency range for fitting the spectra. In conventional data, however, spectral plateaus are regularly present and will be discussed in this section because this can pose a severe challenge for estimating the aperiodic exponent: it shrinks the frequency range at which the 1/f exponent may be examined.

In Fig. 2 a), a simulation of an aperiodic PSD with an exponent of β *=* 2 is shown. By adding white noise, a plateau can be observed starting at 100 Hz in the high-frequency range. Here, we define the onset of the plateau as the lowest frequency of a 50 Hz frequency interval with a vanishing exponent (i.e., flattening of the spectrum). Specifically, we apply FOOOF without periodic peak fitting to measure the slope from 1–50 Hz. We then gradually shift this interval by 1 Hz towards higher frequencies and fit the slope again. We repeat this procedure until the estimated slope reaches a value below β_*thresh*_*=* 0. 05.

**Fig. 2.**
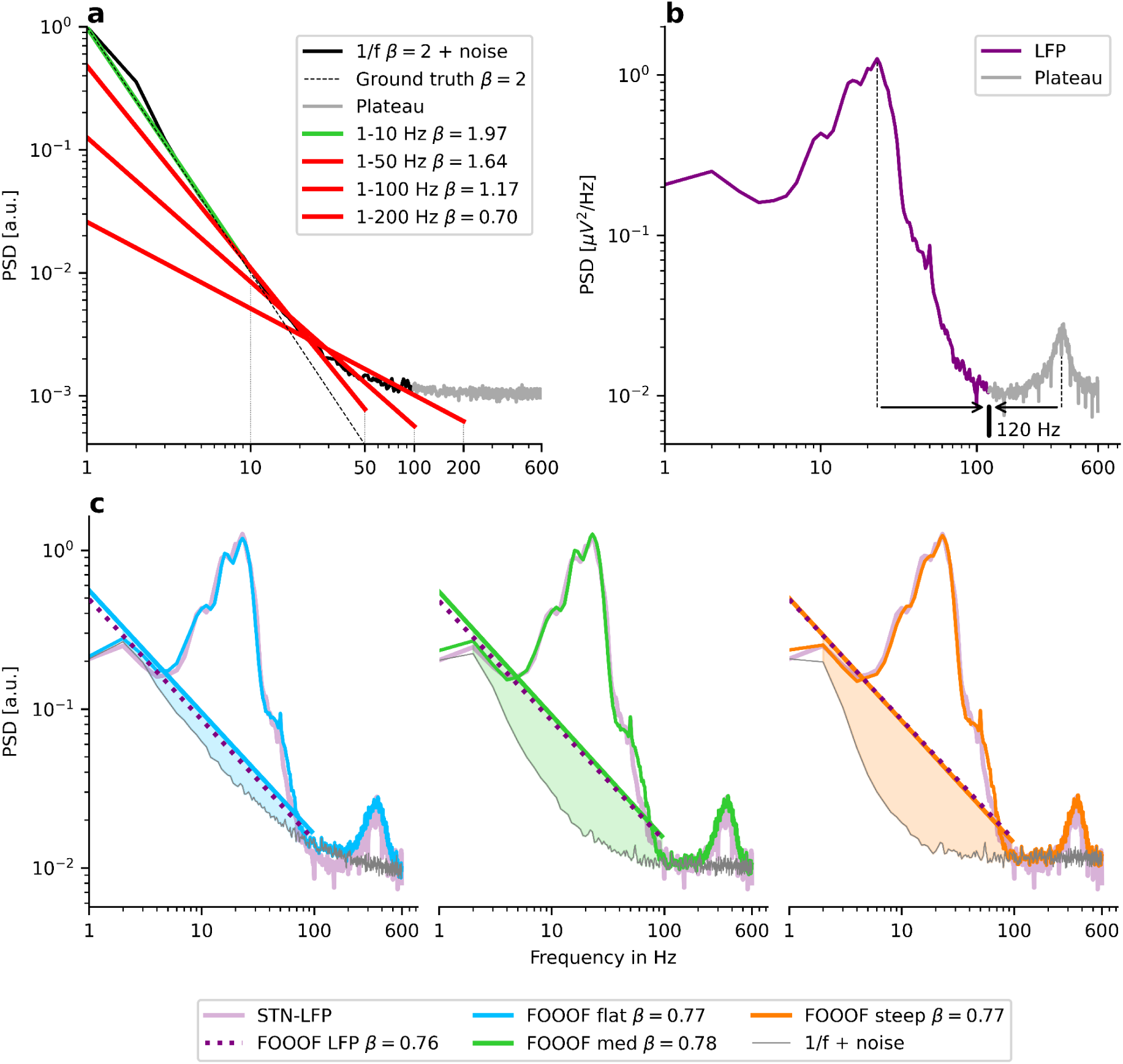
*The spectral plateau disrupts the 1/f power law. The x-axis and the y-axis indicate frequency and PSD, respectively. a) Simulation of an aperiodic PSD (black) with a plateau starting at* 100 *Hz (grey). The spectrum starts to deviate from the ground truth (dashed line) after around* 10 *Hz. Applying FOOOF yields smaller 1/f exponent estimates with larger upper fitting range borders. b) A Parkinsonian LFP spectrum from the subthalamic nucleus shows large oscillations that hinder the plateau onset’s precise detection. c) Adding oscillations of various powers and widths on top of different aperiodic ground truths yields the same 1/f estimation of* β ≈ 0. 77 *in FOOOF. The ground truths are* β = 1 *(blue),* β = 1. 5 *(green), and* β = 2 *(orange)*

We apply FOOOF in the frequency intervals 1−10 Hz, 1−50 Hz, 1−100 Hz, and 1–200 Hz which yields estimated 1/f exponents of β *=* 1. 97, β *=* 1. 64, β *=* 1. 17, and β *=* 0. 70, respectively. The spectral plateau gradually biases the estimated 1/f exponents towards smaller values starting already at 10 Hz. This challenge might be to some extent alleviated if the analysis aims to study differences between groups or experimental conditions such that relative changes of the exponent are most important. However, when the precise onset of the spectral plateau varies across conditions, the upper fitting range border should be chosen as low as possible to minimize this unequal bias. Even if the plateaus seem to be similar across conditions, a lower upper fitting range border will increase the signal-to-noise ratio of the exponent estimates if we define the 1/f ground truth as signal and the impact of the plateau as (possibly Gaussian) noise. We, therefore, recommend choosing low upper fitting range borders and determining the precise onset of the plateau across conditions in order to estimate the exponents under equivalent conditions.

Yet, measuring the onset of the plateau can be difficult in practice if oscillatory peaks mask it. For example, the spectrum in Fig. 2 b) appears to have a spectral plateau onset at 120 Hz. The presence of the high-frequency oscillation peaking at 360 Hz, however, produces a positive slope between 160 Hz and 320 Hz, potentially masking a continued 1/f trend of the spectrum. Accordingly, one cannot exclude the possibility that the actual onset of the plateau is at a higher frequency value (e.g., 200 Hz). On the other hand, the large oscillation ranging from 6 Hz to 100 Hz, peaking in the beta range at 25 Hz, counteracts this effect: In theory, one also cannot exclude that the actual flattening occurs already at 20 Hz.

To demonstrate this effect, for Fig. 2 c) we simulate three power spectra with three different1/f exponent ground truths of β *=* 1 (blue), β *=* 1. 5 (green), and β *=* 2 (orange). Next, we add eight oscillations at 3 Hz, 5 Hz, 10. 5 Hz, 16 Hz, 23 Hz, 42 Hz, 50 Hz, and 360 Hz and tune the oscillation amplitude and width parameters in all three examples to match the recording of Fig. 2 b) (purple). Finally, using FOOOF, we estimate the 1/f exponent in the frequency range from 1–95 Hz in the three simulated and the real PSD. Despite strongly diverging 1/f exponent ground truths, FOOOF estimates an 1/f exponent of about β ≈ 0. 77 in all four cases. The diverging ground truths are apparent in Fig. 2 c) because the true aperiodic component (which is shown in light grey and is invisible to FOOOF) has a plateau onset at high frequencies in the blue curve, at intermediate frequencies in the green, and at low frequencies in the orange curve. However, neither for FOOOF nor for the experimental observer, it is possible to know which of these three scenarios best reflects the real spectrum in Fig. 2 b). Therefore it is difficult to determine at which frequency scale an 1/f estimate might be valid.

Note that this challenge applies not only to cases where the goal is to estimate the 1/f exponent but also when such an estimate is used in order to remove the aperiodic component from the spectrum. While fitting a “shoulder” allows for modeling such a plateau, the strongly varying “shoulder” onsets in Fig. 2 c) cannot be captured, given that the three power spectra share the same appearance. Specifically, the oscillation power estimates based on FOOOF would be almost identical in all three spectra. However, in the simulation, the oscillation power increases considerably from the blue to the green to the orange curve.

### Recommendations

*Scenario A: The power spectra have a plateau onset at higher frequencies, and oscillations do not mask it:*

Challenge 1 does not apply.

*Scenario B: The power spectra have a plateau onset at lower frequencies, and the onset is easily discernible (because no or just small oscillations are present)*.

Determine the precise plateau onsets across conditions. Choose the upper fitting range border as low as possible to increase SNR.

*Scenario C: The power spectra potentially have a plateau onset at lower frequencies, but oscillations mask the exact onset*.

The upper fitting range border must be lower than the onset of the masking oscillation. If the remaining frequency range is too small (as in Fig. 2 b)), aperiodic fitting should be avoided.

### Challenge 2: Avoiding oscillations crossing fitting range borders

When choosing the fitting range to model the aperiodic component, oscillations crossing the fitting range borders must be avoided for all investigated power spectra. FOOOF assumes all oscillation peaks lying within the fitting range because it does not fit partial Gaussian peaks. Consequently, the estimated 1/f exponent error becomes large if the lower or upper fitting range border overlaps with a spectral peak.

In the upper panel of Fig. 3 a), we simulate a PSD with a slope of β = 2 and oscillation peaks at 5 Hz, 15 Hz, and 35 Hz (black graph). We fix the upper fitting range border at 100 Hz and measure the 1/f exponent for all lower fitting ranges from 1–100 Hz up to 80–100 Hz. The lower panel in Fig. 3 a) indicates the absolute error of the estimated slope as a function of the lower fitting range border (red). The error is the absolute deviation from the ground truth |β_*truth*_ − β_*FOOOF*_| Note that the error function resembles the oscillatory peaks, with the greatest errors occurring approximately at the peak center frequencies. The FOOOF parameters are kept at the default setting.

**Fig. 3.**
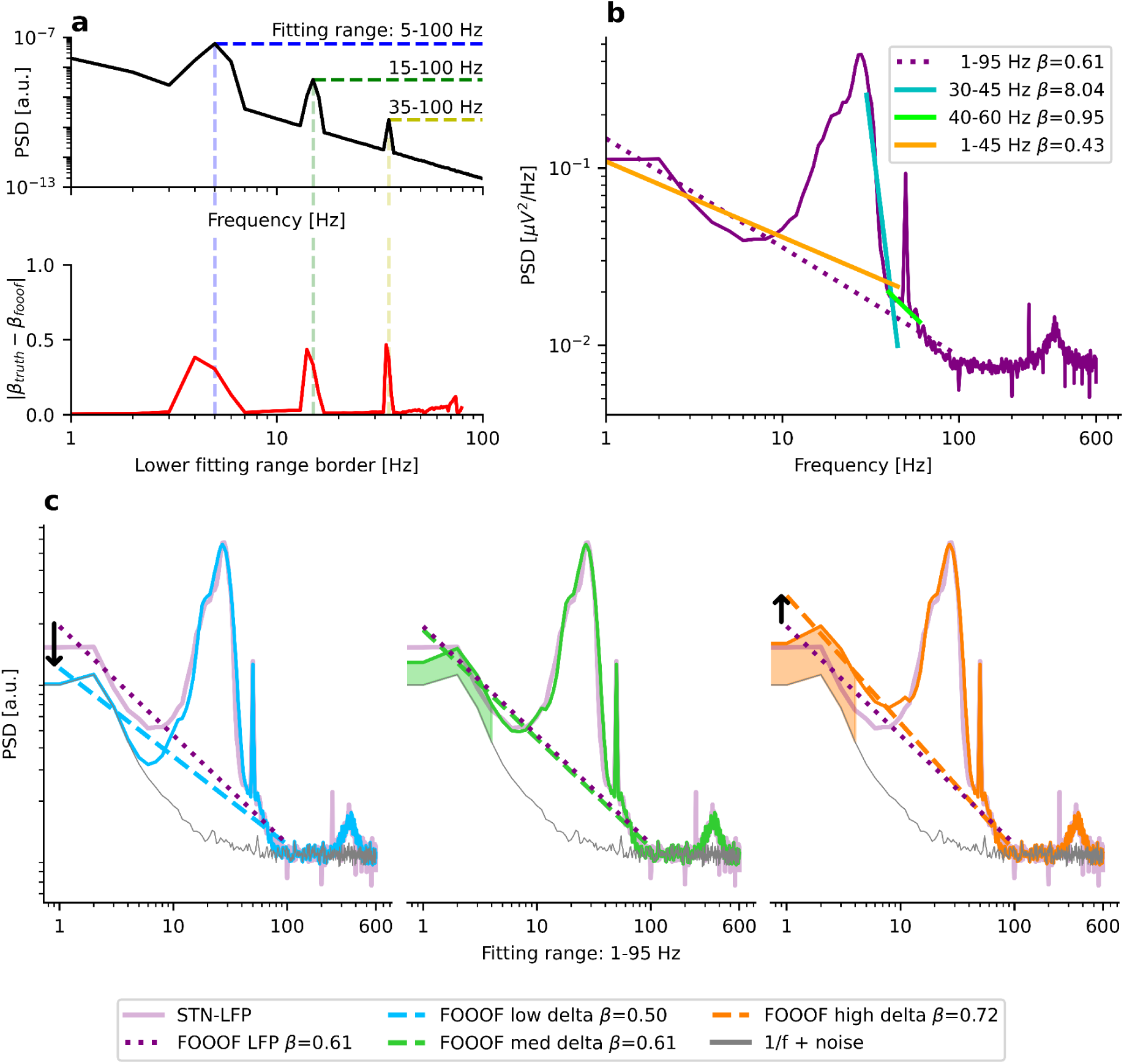
*Oscillations must not cross fitting range borders. a) Upper panel: PSD of a simulated spectrum with β =2 and oscillations at 5 Hz, 15 Hz, and 35 Hz (black). The x-axis and the y-axis indicate frequency and PSD, respectively. Lower panel: The exponent* β *is measured using FOOOF for all 80 frequency ranges from 1–100 Hz to 80–100 Hz (red). The x-axis indicates the lower fitting range border, while the y-axis shows the absolute deviation from the ground truth. b) Various frequency ranges commonly used for E–I estimation are applied to an STN-LFP PSD of a Parkinsonian patient (purple). Since many of the chosen ranges overlap with spectral peaks, the estimated exponents* β *are strongly differing. FOOOF parameters: max_n_peaks=0 (for 30–45 Hz); max_n_peaks=1 (for 40–60 Hz); peak_width_limits=(1, 100) (for 1–45 Hz and 1–95 Hz). c) The simulated PSD in the middle panel (green) was tuned to match the empirical PSD in b) (purple). FOOOF estimates a similar aperiodic exponent for the simulated and the real spectrum (β =0.61). When decreasing the power of the 2 Hz delta oscillation (blue), the estimated aperiodic exponent decreases (β =0.50) despite a constant exponent for the simulated spectrum. When increasing the power of the delta oscillation (orange), the estimated aperiodic exponent increases (β =0.72)*

If the peaks do not completely lie within the fitting range, very error-prone fits are obtained, as shown in another exemplary STN-LFP recording from a Parkinsonian patient (purple) in Fig. 3 b). The fit from 30–45 Hz (turquoise), a frequency range commonly used for estimation of E–I balance, measures the slope of the beta-to-gamma peak, not of the aperiodic component. For this fit, we set the FOOOF parameter for the maximum number of allowed peaks (*max_n_peaks)* to 0 since this frequency range is usually chosen to avoid oscillations altogether. Therefore, FOOOF just fits a straight line without peak modeling. The 40–60 Hz range (green) lies on top of the beta-to-gamma peak, too. Here, we set (*max_n_peaks)* to 1 to account for the power line noise. Further, the 1–45 Hz range (orange) is inappropriate because its upper fitting range border at 45 Hz lies in the middle of the gamma peak. While these are obvious examples of ill-chosen fitting ranges, in practice more subtle (but similar) errors might occur. Therefore the presence of oscillations at fitting range borders must be carefully checked for every single PSD of interest.

The 1–95 Hz range (purple dotted) seems to be the only acceptable range for this spectrum: The upper fitting range border extends beyond the beta-to-gamma peak but ends before the onset of the spectral plateau. The estimated exponent has a value of β_*FOOOF*_ *=* 0. 61. For these two frequency ranges (1–45 Hz and 1–95 Hz), we increased the peak width limits from 0.5–12 Hz (default) to 1–100 Hz to account for the chosen spectral resolution (1 Hz) and enable modeling of the very broad (>12 Hz) beta-to-gamma peak. The corresponding FOOOF fits of Fig. 3 b) are shown in SI Fig. 2.

While the 1–95 Hz range seems best, it appears almost impossible to avoid low-frequency oscillations crossing the lower fitting range border. If some delta oscillations are present, they lead to a steepening of the spectrum which impacts the estimation of the 1/f exponent. We visualize this effect by reproducing the empirical LFP spectrum in three simulations in Fig. 3 c). We set the oscillation frequencies to 2 Hz, 12 Hz, 18 Hz, 27 Hz, 50 Hz (gamma), 50Hz (power line), and 360 Hz. In the panels in Fig. 3 c) from left to right, we only vary the delta power at 2 Hz while keeping the aperiodic component and all other oscillations’ amplitudes and widths fixed. Since the delta oscillation has a bandwidth crossing the lower fitting range border of 1 Hz, FOOOF-estimates of the 1/f exponent diverge strongly between the three scenarios (same FOOOF parameters as for the 1–95 Hz range in Fig. 3 b)). While the aperiodic (white noise-free) ground truth remains unchanged at β *=* 1. 5 for all three simulations, FOOOF estimates an 18% lower 1/f exponent (blue, β *=* 0. 50) if the delta oscillation from the middle panel (green, β *=* 0. 61) is removed. On the other hand, it estimates an 18% larger 1/f exponent (orange, β *=* 0. 72) if we double the power of the delta oscillations. The power of the true delta oscillations in the purple curve is, of course, unknown.

Overall, fitting and removing delta oscillation peaks seems unfeasible since they rarely occur as a single distinguishable peak in the double logarithmic representation. Furthermore, FOOOF requires smooth input spectra to reduce the impact of noise which at the same time hinders fitting sharp peaks. Therefore, we recommend 1/f estimation for a higher lower-border of the fitting range to avoid the impact of these low-frequency oscillations. For high lower borders of the fitting range, oscillations can be better avoided, and if they are present, they likely have less impact on the estimation.

Estimating the power of low-frequency oscillations by removing the aperiodic part of the spectrum poses a special challenge in this regard. Many studies (Donoghue et al., 2020; El Boustani et al., 2009; Fransson et al., 2013; Freeman & Zhai, 2009; Miller et al., 2009; Wen & Liu, 2016) have conceptualized the aperiodic part of the spectrum as self-similar, or fractal, across a wide range of frequencies, such that the estimation of the 1/f exponent should be independent of the chosen fitting range. If this assumption does not hold, however, the aperiodic component must be fitted in the given frequency range of interest. If this frequency range of interest coincides with low-frequency oscillations, this challenge cannot be avoided.

The impact of (sub-)delta oscillations should therefore be kept in mind as a limitation. If one finds a difference of the 1/f exponent between groups of investigation, one should check whether the delta power of the FOOOF-fits varies across conditions. If delta power is similar across conditions but the slope varies, it seems likely that indeed the aperiodic component causes these differences in the estimated slopes and not a distortion by delta oscillations. If delta power does change across conditions (without a global offset of the PSD across all frequencies), the change of slopes could either be caused by a change of oscillatory delta activity (as shown in SI Fig. 1) or by a change in the aperiodic component itself, and these two possibilities cannot be differentiated with full certainty.

### Recommendation

*Scenario A: The 1/f exponent needs to be estimated:*

Use a fitting range at higher frequencies (for example 40–60 Hz) to avoid distortion by low-frequency oscillations.

*Scenario B: The aperiodic component needs to be removed from the PSD:*

If the assumption of self-similarity across a wide range of frequencies holds for the aperiodic part of the spectrum, both slope and intercept of its linear fit could theoretically be obtained from any frequency range. In reality, different exponents could be present in different frequency ranges. In that case, the exponent should be estimated in the broadband range starting at very low frequencies. For this lower fitting range border (starting often at around 1 Hz), the challenge cannot be avoided and should be kept in mind as a potential limitation of the results.

### Challenge 3: FOOOF cannot characterize oscillation peaks that are not clearly distinguishable

As illustrated in Fig. 1, FOOOF models oscillations as Gaussian functions fitted to peaks in the flattened PSD. While this does not impose a severe challenge for clearly isolated peaks, the modeling becomes complicated when peaks overlap partially. If many different peaks overlap, the resulting PSD can be caused by various combinations of oscillations with different frequencies and powers that are impossible to disentangle on a single spectrum. Furthermore, whereas spectral leakage from oscillations at neighboring frequencies but same Fourier phase can add up in different combinations to yield similar power spectra, oscillations at different phases can also subtract power from other peaks.

In the right panel of Fig. 4 a), we present a real PSD that might exemplify a spectrum containing many strongly overlapping oscillation peaks. The underlying time series was recorded from a subject with epilepsy during an absence seizure using a bipolar montage of EEG electrodes F3-C3. Pre-seizure activity is highlighted in turquoise, seizure activity in red, and post-seizure activity in yellow. Absence seizures are proposed to be related to cortico-thalamic E–I dysbalance (Onat et al., 2013) caused by reduced cortical inhibition (Tan et al., 2007), hyperexcitable somatosensory neurons (Karpova et al., 2005), GABA_B_ receptor dysfunctions (Inaba et al., 2009; Merlo et al., 2007), changes in NMDA (D’Arcangelo et al., 2002; Pumain et al., 1992), or mGLU2/3 receptors (Ngomba et al., 2005). It would be interesting to complement these molecular rodent studies by non-invasive human electrophysiological recordings. Specifically, using the 1/f exponent as a biomarker of E–I balance before, during, and after the seizure might help to gain new insights into absence seizures. However, the non-sinusoidal 3 Hz spike-wave discharges might create many harmonic peaks throughout the spectrum. Applying FOOOF (default parameters) in a frequency range of 1–100 Hz yields estimated 1/f exponents of β_*pre*_ *=* 1. 52, β_*seiz*_ *=* 2. 31, and β_*post*_ *=* 1. 52. One could interpret this finding as an increase of the aperiodic 1/f exponent during the seizure, indicating (quite counterintuitively) stronger neural inhibition. However, even though FOOOF subtracts a substantial part of the harmonic peaks by modeling them as four broad peaks with center frequencies at 11 Hz, 22 Hz, 37 Hz, and 50 Hz (see SI Fig. 3), it is not clear whether it can correctly estimate the peak heights. A peak height is the power of an oscillation on top of the aperiodic component. However, there is no reference point for the aperiodic component from which the height could be measured in the scenario of many overlapping oscillations. Hence, it might be that the aperiodic exponent does not change during the absence seizure—instead, the inaccurately removed 3 Hz-harmonics likely caused the increased 1/f exponent value.

**Fig. 4.**
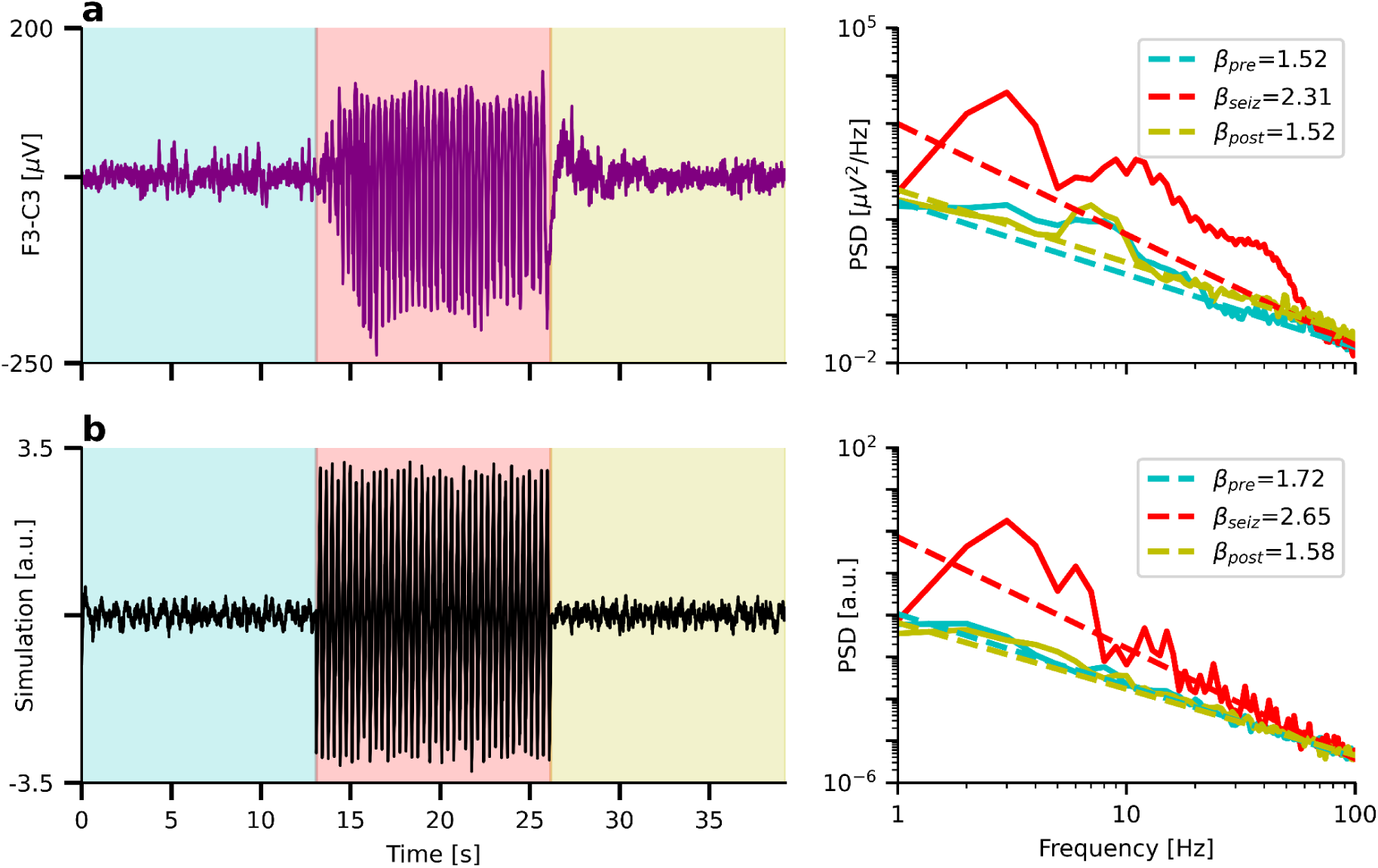
*FOOOF cannot characterize oscillation peaks that are not clearly distinguishable. a) Left: Time series of an absence seizure measured using EEG. Turquoise: Pre-seizure, red: seizure, yellow: post-seizure activity. Right: Corresponding PSDs and aperiodic FOOOF fits. Note the increase of the 1/f exponent during the seizure. b) Left: Simulated 1/f noise and temporarily (red) added 3 Hz saw-tooth signal. Right: Aperiodic FOOOF fits. Note the increase of the 1/f exponent despite constant ground truth of* β*_truth_* = 1. 8

Fig. 4 b) shows a time series of simulated 1/f noise with an exponent of β_*sim*_*=* 1. 8. During the same time interval in which the absence seizure in Fig. 4 a) occurs, we add a saw-tooth oscillation of 3 Hz to the signal. As in the example of the real seizure in Fig. 4 b), FOOOF estimates a strongly increased 1/f exponent even though the ground truth exponent remains constant. The corresponding model (default parameters) is shown in SI Fig. 3.

Note that it is possible to enable FOOOF fitting of the many harmonious peaks by reducing the maximum peak width limits to 1 Hz. While it is not feasible to tune the parameters across conditions (there is an alpha peak with a peak width larger than 1 Hz in the pre-and post-condition), even with the specifically tuned parameters, FOOOF returns increased 1/f exponents β *=* 2. 28 and β_*sim*(*tuned*)_ *=* 1. 91 for real and simulated data, respectively (SI Fig. 3).

### Recommendation

*Scenario A: The PSD appears as a straight line with well-distinguishable peaks on top of this line:*

Challenge 3 does not apply.

*Scenario B: The PSD might contain overlapping peaks:*

The more peaks overlap, the less accurate the model results will be. The lower and upper fitting range borders must vastly extend the overlapping oscillation peaks (challenge 2) to enable peak removal. Estimating the power of overlapping peaks will be difficult.

*Scenario C: Almost the full PSD seems to consist of overlapping peaks (as in Fig 4):*

Avoid fitting the aperiodic component.

## IRASA

### Challenge 1: The evaluated frequency range is larger than the fitting range

While FOOOF tries to iteratively fit all oscillatory peaks to obtain a periodic model, IRASA takes the median of spectra after up- and down-sampling to eliminate the peaks, as shown in Fig. 1. As a result, it aims to obtain the pure aperiodic component that is assumed to be invariant to resampling. As an advantage over FOOOF, IRASA can overcome challenge 2: Even if a peak crosses the fitting range border (at the original sampling rate), it can be removed successfully due to the resampling procedure.

In Fig. 5 a), we replot the spectrum of Fig. 3 a) and estimate the 1/f exponent for all frequency ranges between 1–100 Hz and 80–100 Hz. In contrast to FOOOF, IRASA has minimal errors for all frequency ranges. The reason is that the fitting range of FOOOF has well-defined borders: If the lower border is set to 5 Hz (the center frequency of the first peak), it cannot identify and model the 5 Hz peak correctly. On the other hand, for IRASA, the fitting range is blurry: By up- and down-sampling the spectrum, the peaks are shifted towards lower and higher frequencies. Therefore, the evaluated frequency range of IRASA is much more extensive than the actual fitting range.

**Fig. 5.**
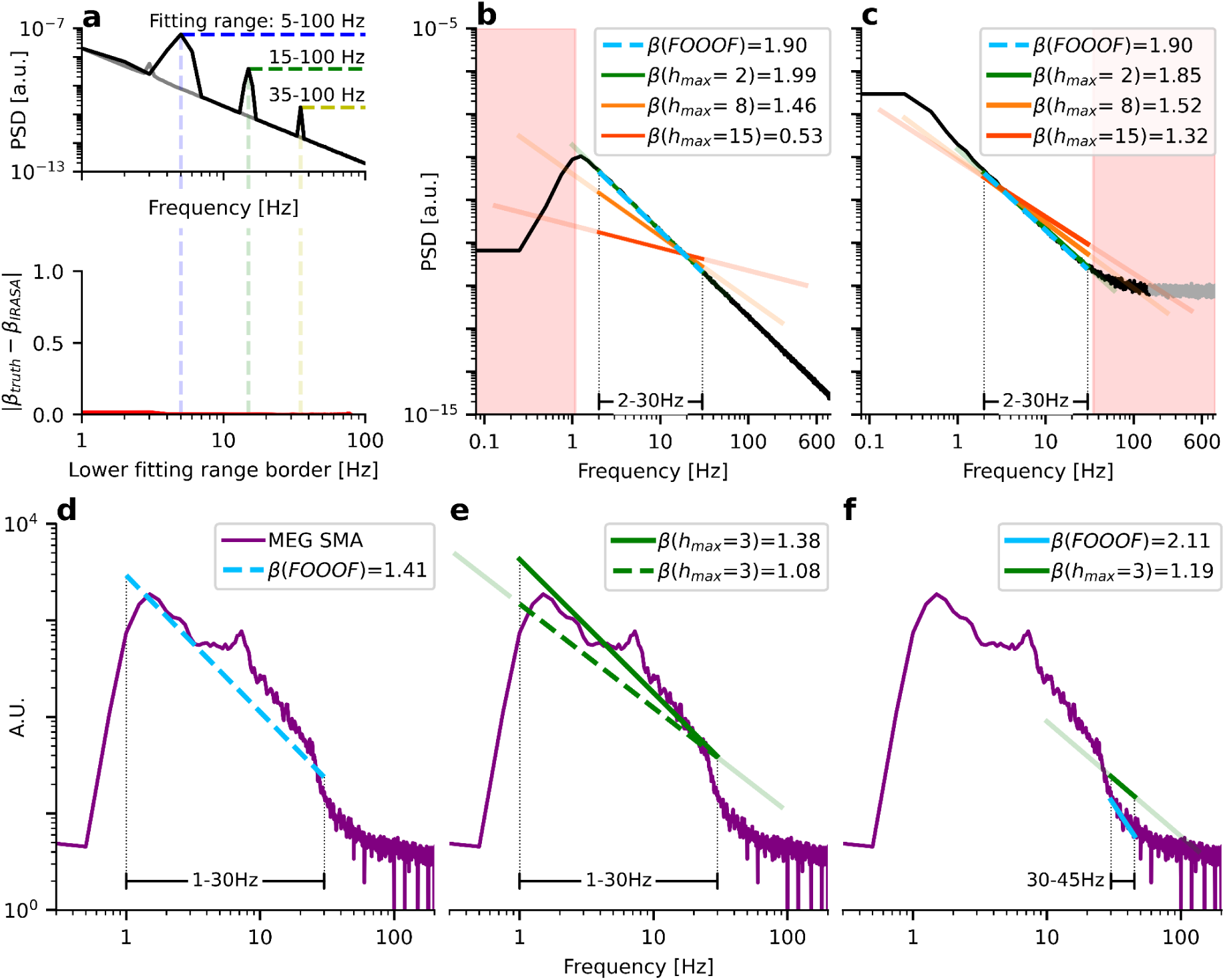
*IRASA’s evaluated frequency range is larger than the fitting range. a) Upper panel: Same simulation as in Fig. 3 a). Lower panel: The lower fitting range border is shown on the x-axis, the absolute deviation from the ground truth on the y-axis. IRASA correctly estimates the 1/f exponent for all used fitting ranges. b) Simulated aperiodic PSD with a ground truth of* β = 2. *A 1 Hz highpass filter disrupts the 1/f power law. IRASA’s fitting range for the maximum resampling factor h_max_* ∈ {2, 8, 15} *is indicated as bright-colored lines upon the fitted aperiodic components, with the evaluated frequency ranges after up- and down-sampling indicated in corresponding transparent colors. IRASA’s error of the 1/f estimation increases with larger resampling rates h_max_ (and lower resampling rates 1/h_max_, respectively). c) Same as b) with a spectral plateau disrupting the 1/f power law. d) FOOOF 1/f estimate within 1–30 Hz for a spectrum obtained from voxel data after MEG source reconstruction. e) IRASA 1/f estimates for an evaluated frequency range of 1–30 Hz (green) and an evaluated frequency range of 0.3–90 Hz (green-dashed, corresponding to a fitting range of 1-30 Hz at h_max_* = 3*). f) FOOOF (blue) and IRASA (green) estimates of the 1/f exponent for the same fitting range of 1–30 Hz*

For example, if we chose only two resampling factors *h*_*set*_ *=* {2, 3}, the spectrum would be up-sampled by *h*_*up*1_ *=* 2 and *h*_*up*2_ *=* 3 and down-sampled by *h*_*down*1_*=* 1/2 and *h*_*down*2_ *=* 1/3. As a result, a fitting range of 10–100 Hz would correspond to four evaluated frequency ranges of 20–200 Hz, 30–300 Hz, 5–50 Hz, and 3.3–33 Hz. Of these four resampled spectra, IRASA takes the median. The lower border of the evaluated frequency range *f*_*eval. min*_ can be calculated from the minimum fitting range border *f*_*fit min*_ divided by the maximum resampling factor *h*_*max*_ according to equation 1. The upper evaluated frequency border *f*_*eval. max*_ corresponds to the upper fitting range border *f*_*fit max*_ multiplied by *h*_*max*_ according to equation 2.

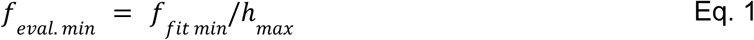

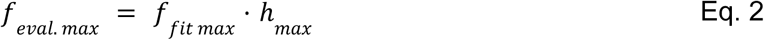

While evaluating a larger frequency range than the actual fitting range can be advantageous, as shown in Fig. 5 a), it can also lead to severe challenges, as shown in Fig. 5 b). Here, we simulate an aperiodic PSD with β *=* 2, which is highpass filtered at 1 Hz. We then apply IRASA in a fitting range of 2–30 Hz for three different h-sets with maximum resampling factors *h*_*max*_ *=* 2, *h*_*max*_ *=* 8, and *h*_*max*_ *=* 15, respectively. The fitting ranges, indicated in green, orange, and red, are the same, but the evaluated frequency ranges, shown in the corresponding transparent colors, increase with increasing *h*_*max*_.

Note that the highpass filter disrupts the 1/f power law for low frequencies and violates IRASA’s assumption of a resampling-invariant aperiodic component. With increasing *h*_*max*_, IRASA evaluates substantially larger parts of the low-frequency stopband which increasingly biases its 1/f estimates towards smaller values. A good agreement with the ground truth of β *=* 2 is only obtained for *h*_*max*_ *=* 2, which corresponds to an evaluated frequency range of 1–60 Hz, avoiding the stopband of the highpass-filtered spectrum.

Apart from low-frequency fitting artifacts due to highpass filtering, care must also be taken to avoid fitting artifacts at high frequencies. For example, in Fig. 5 c), the high-frequency spectral plateau disrupts the 1/f power law. Even though the upper fitting range border of IRASA is set well below the plateau onset to *f*_*fit max*_*=* 30 Hz, IRASA does nevertheless evaluate the plateau due to the upsampling step. Therefore, with growing *h*_*max*_, IRASA biases the 1/f estimates towards smaller values again.

Even in the absence of a spectral plateau, care must be taken to avoid the resampled Nyquist frequency. For example, for a sampling rate of *f*_*sample*_ *=* 2400 Hz and *h*_*max*_*=* 10, the resampled Nyquist frequency reduces from *f*_*Nyquist*_ *=* 1200 Hz to *f*_*Nyquist resampled*_*=* 120 Hz. Accordingly, the upper fitting range border must not exceed this value. The same holds true for a potentially applied lowpass filter. In general, to avoid accidentally fitting spectra above Nyquist frequency or in the stopbands of lowpass or highpass filters, it is advisable to choose *h*_*max*_ as small as possible. Furthermore, the evaluated frequency range should always be checked by calculation from the fitting range and *h*_*max*_.

Given that IRASA evaluates a more extensive frequency range than the fitting range, the meaning of the fitting range becomes imprecise. For example, if we are interested in fitting the 1/f exponent from 1–30 Hz and use *h*_*max*_*=* 2, we should choose 2–15 Hz as a fitting range for the IRASA algorithm. However, since only the minimum and maximum resampled spectra contain the 1 Hz and 30 Hz borders of interest, IRASA emphasizes the estimation of the 1/f exponent from intermediate frequency values above 1 Hz and below 30 Hz.Therefore, 1/f exponents estimated by IRASA cannot be directly compared to 1/f exponents estimated by FOOOF.

We visualize this effect for a spectrum of voxel data obtained by MEG source reconstruction in the lower panels d)–f) of Fig. 5. In d), FOOOF estimates an 1/f exponent of β_*FOOOF*_ *=* 1. 41 in the fitting range of 1–30 Hz. Due to the highpass filter, IRASA obtains a lower value of β_*IRASA*_ *=* 1. 09 for the same fitting range which, however, actually corresponds to an evaluated frequency range of 0.33–90 Hz at *h*_*max*_ *=* 3 (Fig. 5 e). Hence, setting the evaluated frequency range to 1–30 Hz (by setting the fitting range to 3–10 Hz) yields β_*IRASA*_*=* 1. 38 which is similar to the FOOOF estimate.

Matching the evaluated frequency range of IRASA to the fitting range of FOOOF is not always possible, though. Consider, for example, the fitting range of 30–45 Hz shown in Fig. 5 e). At *h*_*max*_*=* 3, the evaluated frequency range of IRASA is 10–135 Hz. Due to the spectral plateau, IRASA estimates a much smaller exponent of β_*IRASA*_*=* 1. 22 compared to β_*FOOOF*_*=* 2. 11. This time, we cannot shrink IRASA’s fitting range to match its evaluated frequency range with FOOOF’s fitting range. At *h*_*max*_*=* 3, the lower fitting range border of IRASA must be 3 · 30 Hz *=* 90 Hz to match the lower fitting range border of FOOOF at 30 Hz. However, the upper fitting range border needs to be 45 Hz / 3 *=* 15 Hz to match the upper fitting range border of FOOOF. This would lead to an inverse fitting range of 90–15 Hz. Here, it cannot be avoided that IRASA evaluates a much more extensive frequency range than 30–45 Hz. As a result, FOOOF and IRASA cannot yield comparable 1/f estimates for this frequency range.

### Recommendations

Always calculate the evaluated frequency range from the fitting range and *h*_*max*_ according to equations 1 and 2. Choose the maximum resampling factor *h*_*max*_ as small as possible in order to 1) avoid fitting artifacts, 2) to improve comparability with other methods, and 3) to improve the interpretability of the investigated frequency range.

Set the evaluated frequency range—and not the fitting range—to the frequency range of interest.

### Challenge 2: Broad peak widths require large resampling factors

In challenge 1, we recommend choosing the maximum resampling factor *h*_*max*_ as small as possible. However, for IRASA to work correctly, the resampling factors must be sufficiently large. This is because IRASA shifts the peaks in the frequency scale up and down through up- and downsampling. Therefore, a single peak appears multiple times on the frequency scale (Fig. 1 b). For a range of sufficiently large (and small) resampling factors, the resampled peaks are completely separated and, by taking the median of their geometric mean, subsequently eliminated. However, if the range of resampling factors is too small or the peaks too broad, the resampled peaks overlap. In that case, peak removal by taking the median will not be successful.

In Fig. 6 a), we replot Fig. 5 a). However, by increasing the peak widths from the left to the right panels, the 1/f estimation error of IRASA increases strongly. This is because the peaks cannot be fully separated. As a result, IRASA’s calculated aperiodic component, shown in grey, still contains the up and downsampled peaks after taking the median. Note that not the peak width Δ*f* itself must be sufficiently small to get separated, but instead, Δ*f*_*log*_, as it appears in the logarithmic frequency scale, the logarithmic peak width needs to be sufficiently small. For this reason, a peak width of 4 Hz at a center frequency of 5 Hz has a similar effect as a peak width of 12 Hz at a center frequency of 35 Hz.

**Fig. 6.**
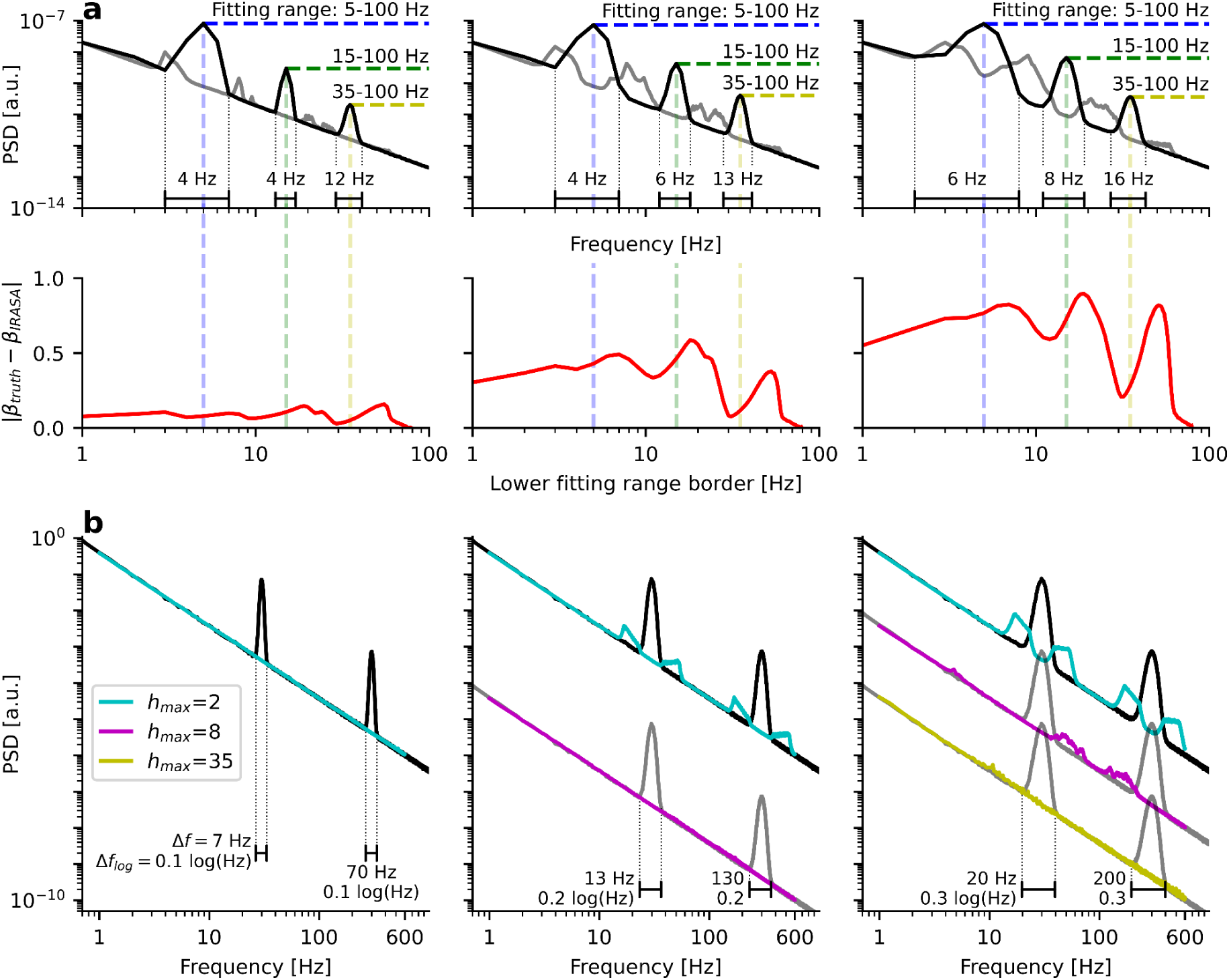
*Broad peak widths require large resampling factors. a) Upper panel: Similar as in Fig. 5 a) but with increasing peak widths from left to right. Note that removal of peaks from the aperiodic component (grey) worsens with broader peak widths. Lower panel: The lower fitting range border is on the x-axis, the absolute deviation from the ground truth on the y-axis. The 1/f exponent estimation error increases with larger peak widths. b) Simulation of a 30 Hz and 300 Hz peak with increasing peak widths from left to right. Larger peak widths require larger resampling factors. Note that not the absolute peak width but rather the logarithmic peak width* Δ*f_log_ determines the minimum resampling factors*

We visualize this effect in panel Fig. 6 b) by simulating a PSD with two oscillations at *f*_1_ *=* 30 Hz and *f*_2_ *=* 300 Hz. The peak width of the second peak is 70 Hz and therefore 10 times as large as the peak width of the first peak. However, on the logarithmic frequency axis, they appear with the same width. A maximum resampling factor of *h*_*max*_ *=* 2 is sufficient to remove the peaks correctly. Thus, they are fully eliminated from the aperiodic component shown in turquoise. However, when the peak widths are increased to a logarithmic value of 0.2 log(Hz), *h*_*max*_ *=* 2 is not sufficient anymore: The up- and down-sampled peaks remain visible in the estimate of the aperiodic component. If we increase *h*_*max*_ to a value of 8, however, peak removal works well. For a further increase of the logarithmic peak width to 0.3 log(Hz), however, *h*_*max*_ *=* 35 is necessary. We visualize this challenge on empirical data of MEG and LFP data of dataset 3 in SI Fig. 4.

We calculated the logarithmic peak width as Eq. 1 Δ*f*_*log*_ *= log*_10_ (*f* _2_ /*f*_1_) where *f*_1_ corresponds to the lower bound of the peak and *f*_2_ to the upper bound of the peak. The bounds were found by calculating the first bin of the PSD, which deviates above a threshold of 0.001 from the aperiodic ground truth. Note that there is no exact equation/heuristic to calculate the minimum *h*_*max*_ as a function of peak width because always many resampling factors *h* are calculated, which will lead to a gradual peak removal depending on the degree of peak separation.

### Recommendations

Choose *h*_*max*_ as small as possible (challenge 1) while keeping it large enough to obtain peak-free estimates of the aperiodic component (challenge 2). If the peaks are very broad and *h*_*max*_ cannot be chosen sufficiently large without avoiding challenge 2, IRASA cannot be applied.

### Challenge 3: IRASA cannot characterize oscillation peaks that are not clearly distinguishable

Similar to FOOOF, IRASA cannot separate strongly overlapping peaks. However, as shown in Fig. 7 b), IRASA performs quite well for dataset 2 because the harmonic peaks do not strongly overlap above 10 Hz. Instead, many local power minima in between the harmonic peaks are very close to the power of the aperiodic ground truth. As a consequence, adding the 3 Hz sawtooth signal only slightly increases the estimated 1/f exponent from β_*pre*/*post*_ *=* 2. 24 to β_*seiz*_ *=* 2. 46. In the middle panel of SI Fig. 5 b), the extracted oscillatory component of IRASA is shown in orange, indicating a good extraction of harmonic peaks at multiple integers of 3 Hz.

**Fig. 7.**
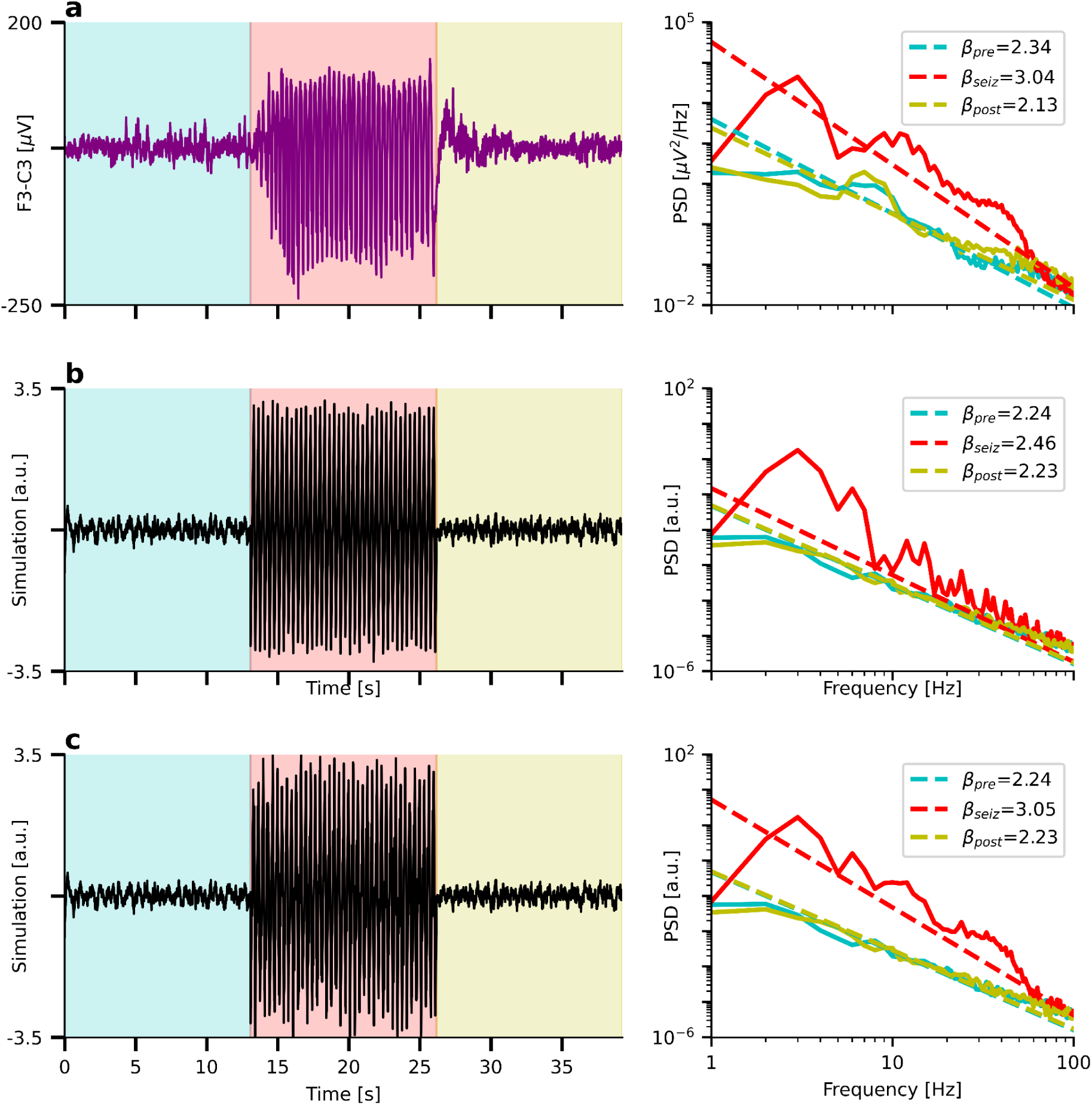
*IRASA cannot characterize oscillation peaks that are not clearly distinguishable. a) and b) left panel: Same as Fig. 4. a) and b) Right panel: Same as Fig. 4 but showing the 1/f fits by IRASA. c) IRASA’s performance on the simulation drops significantly if two strongly overlapping peaks in the alpha (10 Hz) and beta range (25 Hz) are added. Ground truth:* β*_truth_* = 1. 8

If, however, we now add two strongly overlapping oscillations at 10 Hz and 25 Hz, IRASA is no longer capable of successfully removing the peaks. As a result, it now estimates an exponent of β_*seiz*_*=* 3. 05—much larger than the ground truth at β_*truth*_*=* 1. 8.

## Discussion

Both periodic and aperiodic components of power spectra are frequent targets of investigation in electrophysiological studies. The separation of both components before analysis helps to disentangle their relative contribution to the spectrum. FOOOF and IRASA are commonly used for this purpose. It should be highlighted, though, that the methods follow different concepts: Whereas FOOOF models periodic components and a single aperiodic component and outputs the corresponding parameters, IRASA only separates them, allowing further independent processing. Other methods to separate periodic and aperiodic PSD components exist too, for example, eBOSC (Kosciessa et al., 2020).

However, they cannot overcome the method-unspecific challenges in electrophysiological PSDs such as 1) spectral plateau onsets at relevant frequencies, 2) hidden low-frequency oscillations, and 3) overlapping peaks.

Here, we evaluated common challenges of the separation procedure based on two popular methods and summarized both general and method-specific challenges. These challenges apply to EEG, MEG, and LFP data obtained by independent research groups, indicating the general applicability of the results.

### Neurophysiological interpretation

#### Spectral Plateau

The spectral plateau can hinder a correct separation of the PSD components. If the fitting range of FOOOF or the evaluated frequency range of IRASA includes a spectral plateau, the 1/f exponent will be estimated too low. In the presence of periodic components at the spectral flattening, a faulty aperiodic power estimation will lead to a faulty periodic power estimation. In addition, large periodic components could hide the onset of the spectral plateau. This hinders a proper decision on the choice of the upper fitting range border. If the 1/f exponent is estimated too low due to the spectral flattening, this could be misinterpreted as an increased E–I ratio since typically flatter spectra are associated with more pronounced excitability (Gao et al., 2017).

E–I balance estimation is usually applied to estimate relative 1/f differences between conditions. If the spectral plateau onset were to occur at exactly the same frequency for all spectra, a relative 1/f comparison would still be viable. A random fluctuation of the onset would introduce noise to the estimates, and a systematic difference of the plateau onset between conditions would lead to type 1 errors.

The origin of the spectral plateau at high frequency is likely rooted in Gaussian properties of amplifier noise and impedance noise (Scheer et al., 2006; Waterstraat et al., 2015). (Waterstraat et al., 2021) showed that the onset of the plateau starts at higher frequencies if recordings are done with a low-noise MEG system. In addition to system noise, biological high-frequency noise caused by electromyography (EMG) from the head muscles can contribute to spectral flattening, although EMG does not necessarily have a flat spectrum (Muthukumaraswamy, 2013). However, even when recording LFPs from the subthalamic nucleus using a low-noise amplifier, which can be considered as hardly affected by EMG activity, a spectral plateau could be observed one order of magnitude above the system’s noise level (unpublished data). Neuronal population spiking activity probably contributes to this spectral plateau (Belluscio et al., 2012; Buzsáki et al., 2012; Zanos et al., 2011). Better understanding the origins of the spectral plateau is of major interest and requires further research. If it is caused by noise such as systems noise or EMG, an identification of the origin could help to clean the data from this high-frequency plateau. If it has a neurophysiological origin, a thorough analysis of the plateau might yield novel neurophysiological insights.

#### Fitting ranges

The choice of the fitting range depends on the goal of the study and the properties of the data. In the literature, the 1/f exponent was investigated for different frequency ranges such as 0.01–0.1 Hz (He et al., 2010), 0.5–35 Hz (Miskovic et al., 2019), 1–10 Hz (Schaworonkow & Voytek, 2021), 1–20 Hz (Bédard et al., 2006), 1–30 Hz (Wen & Liu, 2016), 1–40 Hz (Colombo et al., 2019), 1–20 and 20–40 Hz (Colombo et al., 2019), 1–100 Hz (He et al., 2010), 2–24 Hz (Voytek et al., 2015), 3–30 Hz (Pereda et al., 1998), 3–55 Hz (Waschke et al., 2021), 10–100 Hz (Freeman & Zhai, 2009), 20–65 Hz (Bédard et al., 2006), 30–50 Hz (Gao et al., 2017; Lendner et al., 2020; Stolk et al., 2019) and 40–60 Hz (Gao et al., 2017)).

If a study using FOOOF aims to generally estimate an 1/f exponent and is free to choose any fitting range for that purpose, we generally recommend avoiding low lower fitting range borders. It is unknown how hidden low-frequency oscillations at the lower fitting range border might impact the 1/f estimate. For example, if the 1/f exponent is compared between two conditions and in one condition there are larger delta oscillations, a fitting range starting from 1 Hz could have a larger y-intercept due to the presence of low-frequency oscillations. This could lead to a larger 1/f exponent and could be misinterpreted as stronger neural inhibition. The same holds for any other lower fitting range border. Therefore, the lower border should be chosen to best avoid known oscillation frequencies, depending on the study.

The same problem applies to IRASA but less severely since this method is not based on a single frequency range but rather, due to up- and downsampling, to a set of different frequency ranges. This comes at the cost of only vaguely defined upper and lower fitting range borders, hindering an easy comparison with fitting ranges used in other studies.

In general, there is no one-range-fits-all fitting range applicable to all kinds of PSDs. Therefore, we recommend examining the PSDs of interest carefully and choosing the fitting range that best avoids the challenges discussed so far in addition to further possible data-specific or goal-specific challenges.

Finally, if the purpose of the 1/f estimation is not to obtain the 1/f exponent but rather the removal of the aperiodic component for better periodic power assessment, a broadband range (such as 1–100 Hz) should be chosen.

#### Overlapping peaks

The stronger periodic components overlap, the more difficult estimating their power becomes. In the shown exemplary data, overlapping peaks occurred mainly in STN data and in dataset 2. MEG and EEG cortical data of healthy participants typically have peaks in the ranges 8–13 Hz and 18–25 Hz which does not impose considerable challenges for the estimation of the aperiodic part of the spectrum. Assessing the 1/f exponent is still feasible if the overlapping periodic components make up only a minor part of the fitting range. On the other hand, if the overlapping peaks make up a majority of the frequency range to investigate, as in Figs. 4, 7, and 8 b), a separation of the periodic and aperiodic components is not recommended and will likely lead to imprecise results. In the case of the absence seizure shown in Figs. 4 and 7, neural inhibition during the seizure is likely overestimated due to overlapping peaks leading to a false 1/f estimation.

**Fig. 8.**
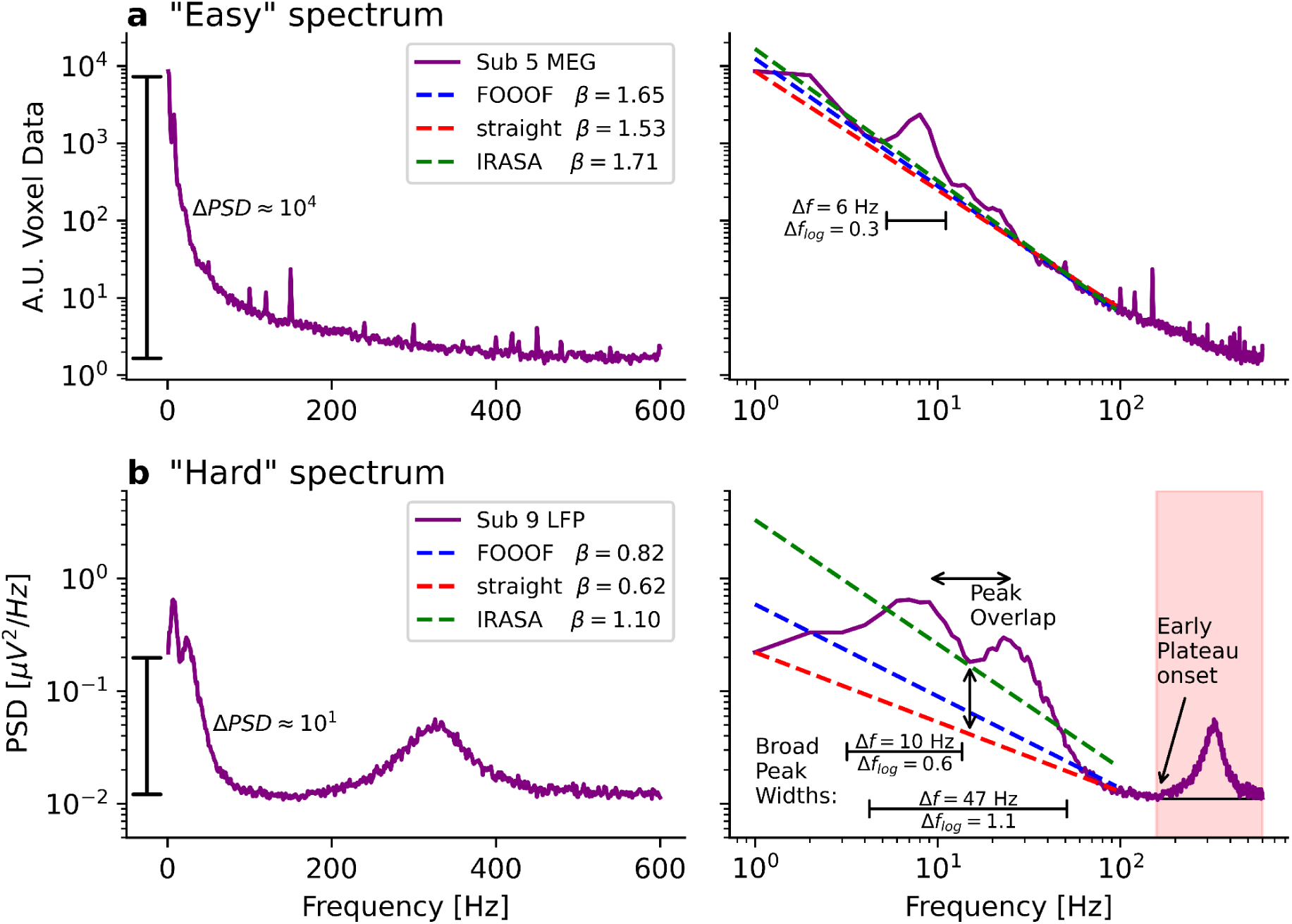
“Easy” and “hard” PSDs. a) Left: Voxel MEG PSD of a Parkinsonian patient on a semilogarithmic scale. Right: Same PSD on a double logarithmic scale. FOOOF, IRASA, and simply connecting the PSD value at 1 Hz to the PSD value at 95 Hz as a straight line (“straight”) yield similar 1/f exponents. We regard such a PSD as “easy” because it avoids all the discussed challenges. b) LFP data of a Parkinsonian patient on a semilogarithmic scale. Right: Same PSD on a double logarithmic scale. FOOOF, IRASA, and “straight” yield different 1/f exponents. We regard such a spectrum as “hard” because it contains many challenges

#### Broad peak widths

In contrast to FOOOF, IRASA cannot handle very broad peaks well. This limitation is especially severe for the analysis of LFPs of Parkinsonian patients (datasets 1 and 3). In the original article (Wen & Liu, 2016), IRASA was only evaluated on pure sine oscillations, for which the method works very well. We, therefore, do not recommend using IRASA if the peaks seem to have broad logarithmic peak widths Δ*f*_*log*_. It is not possible to give a threshold value for a maximum logarithmic peak width as IRASA is, in theory, able to fit any peaks if the *h*-values are chosen sufficiently large. However, in practice, the *h*-values must not exceed a certain range to avoid too low (highpass, Fig. 5 b)) or too high (spectral plateau, Fig. 5 c)) fitting ranges, calculated using Eq. 1 and 2.

#### Estimating E–I balance

(Gao et al., 2017) proposed to use the 1/f exponent as an indicator of E–I balance. Subsequent studies indicated the usefulness of this idea also for non-invasive EEG/MEG data (Colombo et al., 2019; Gao et al., 2017; Lendner et al., 2020; Miskovic et al., 2019; Waschke et al., 2021). While we outlined that 1/f exponent estimation is affected by many possible error sources, we do not argue that it should be avoided altogether. While proper 1/f estimation seems to be beyond reach for some PSDs (for example, the one shown in Fig. 8 b)), it seems to be a promising measure for others (Fig. 8 a)). Thus we suggest that existing methods could be enhanced by more elaborate data cleaning, such as spatio-spectral decomposition (SSD) (Nikulin et al., 2011), independent component analysis (ICA), or inverse modeling. Moreover, it might be possible to develop new methods that measure the 1/f exponent more reliably than the ones discussed in the present study. For example, if the periodic and aperiodic components are assumed to vary over time independently, it could be possible to disentangle them using machine learning algorithms such as non-negative matrix factorization (Lee & Seung, 1999). And it might become possible to measure E–I balance through other electrophysiological measures thus further validating the 1/f exponent of the PSD. We elaborate on this below.

(Bruining et al., 2020), for example, proposed to measure E–I balance based on the alpha band amplitude envelope and its detrended fluctuation analysis (DFA) exponent (Peng et al., 1995). (Stephani et al., 2020) related the N20 of somatosensory evoked potentials to cortical excitability. Other researchers related spontaneous fluctuations of alpha-band power to E–I balance (Romei et al., 2008) and (Iemi et al., 2019) found alpha- and beta-band power to predict suppression of ERP-components, which was interpreted as increased inhibition. This relationship held true even after controlling for fluctuations in the 1/f exponent. It might be possible to estimate E–I balance by measuring transcranial magnetic stimulation (TMS) evoked potentials using EEG. By combining these two methods, (Massimini et al., 2005) showed a breakdown of effective cortical connectivity during non-rapid eye movement (REM) sleep. Effective connectivity was also related to the 1/f exponent by (El Boustani et al., 2009). The perturbational complexity index (Casali et al., 2013) follows these lines to separate unconscious states of low excitability (non-REM sleep, anesthesia) from conscious states of high excitability (wakefulness, REM sleep). Indeed, (Colombo et al., 2019) could link this index to the 1/f exponent during wakefulness and anesthesia yielding similar results with both methods. However, it should be noted that REM sleep (a conscious state of mind) is associated with a larger 1/f exponent compared to NREM sleep (unconscious) while NREM sleep is associated with a larger 1/f exponent compared to wakefulness (conscious) (Lendner et al., 2020). These findings agree with in vivo calcium imaging measurements of E–I balance in mice during wakefulness, NREM sleep, and REM sleep (Niethard et al., 2016).

In the best scenario, different methods used for E–I estimation will lead to similar results and might be used in conjunction. So far, the relationship between 1/f exponent and E–I balance remains a hypothesis to be further validated.

### Computational cost and parameter tuning

From a computational perspective, FOOOF is much faster than IRASA. When applied to 9 time series of dataset 1 (ca. 180 s at *f*_*sample*_ *=* 2400 Hz corresponding to ≈ 9 × 440, 000 data points), parameterization with FOOOF was about 50 times faster than separation with IRASA when the PSD calculation time was included. FOOOF was 100 times faster if the PSDs were precalculated. For the comparison, we used 7 runs and fitted a frequency range from 1–30 Hz. For FOOOF, the default parameters were chosen, and for IRASA a window length of 4 s and a set of 17 resampling factors *h*_*set*_ *=* {1. 1, 1. 15, …, 1. 9} was used. FOOOF computation slows down when the PSDs have a very high resolution leading to many iterations of fitting noise peaks. IRASA computation slows down when the number of resampling factors is increased and when their values are increased.

Increasing IRASA’s resampling values can help with very broad peak widths (challenge 2) but simultaneously enlarges the evaluated frequency range (challenge 1). Increasing the number of resampling factors beyond 17 or changing the window length does not help with the challenges presented in this article. FOOOF requires extensive parameter tuning for optimum results, but the posed challenges cannot be resolved by parameter selection. In general, the fitting range of FOOOF and the evaluated frequency range of IRASA are the most critical parameters for each method.

## Conclusion

To study either periodic or aperiodic PSD components, it is useful to disentangle both components. As there are theoretically infinite solutions to this inverse problem, it is probably neither possible to perfectly separate them nor to evaluate and verify a performed separation since the ground truth remains unknown. Some PSDs seem to be particularly easy to separate because they avoid most of the discussed challenges. For those PSDs, we generally recommend performing a separation to study the periodic or aperiodic components in a more isolated manner. We give an example of such an “easy” PSD in Fig. 8 a).

These “easy” PSDs appear as an almost straight line in double logarithmic space with some well-distinguishable, narrow periodic peaks on top of it. There is no spectral plateau disrupting the 1/f power law and the y-axis of the PSD extends over 4 orders of magnitude. When applying FOOOF and IRASA from 1–95 Hz or simply connecting values at 1 Hz and 95 Hz to a straight line in double logarithmic space, similar values (β_*FOOOF*_*=* 1. 65, β_*IRASA*_*=* 1. 71, β_*straight*_*=* 1. 53) are obtained for the 1/f exponent.

Other PSDs seem to be very difficult to separate. For such, we recommend avoiding the separation since the results will be arbitrary and might lead to ill-informed interpretations. An example of such a “hard” PSD is shown in Fig. 8 b). These spectra do not appear as a straight line. They have very broad and overlapping peaks and a spectral plateau onset at lower frequencies. As a result of this plateau, the y-axis spans only one order of magnitude. When applying FOOOF, IRASA, or a straight-line connection between 1 Hz and 95 Hz, strongly diverging 1/f exponent values (β_*FOOOF*_*=* 0. 82, β_*IRASA*_*=* 1. 10, β_*straight*_*=* 0. 62) are obtained.

Checking PSDs for the challenges discussed in this work will help to decide whether a technique to separate neuronal oscillations from aperiodic 1/f activity should be applied, which algorithm to use, and which parameters to choose.

## Declarations

## Acknowledgments

This work was supported by Deutsche Forschungsgemeinschaft (German Research Foundation) Project ID 424778381 TRR 295. The Wellcome Centre for Human Neuroimaging is supported by core funding from Wellcome [203147/Z/16/Z]. VL is grateful to the National Hospital for Neurology and Neurosurgery clinical team for their help with data collection.

## Conflict of interest

The authors have no conflicts of interest to declare that are relevant to the content of this article.

## Data availability

The datasets analyzed in the present study are not publicly available due to privacy regulations of patient health information but are available from the corresponding author upon reasonable request.

## Code availability

The code of the entire study is available at https://github.com/moritz-gerster/separating_periodic_from_aperiodic_PSDs

## Author’s contributions

Conceptualization: MG, GW, VN, GC; Methodology: MG, GW, VN, GC; Data curation: VL, KL, EF, AS; Formal analysis and investigation: MG; Visualization: MG; Writing - original draft preparation: MG; Writing - review and editing: MG, GW, VL, KL, AS, EF, GC, VN; Funding acquisition: VN, GC; Resources: Not applicable; Supervision: VN, GW, GC.

## Ethics approval

Please refer to the methods section.

## Consent to participate

Please refer to the methods section.

## Consent for publication

Please refer to the methods section.

## Supplementary Information

**SI Fig. 1.**
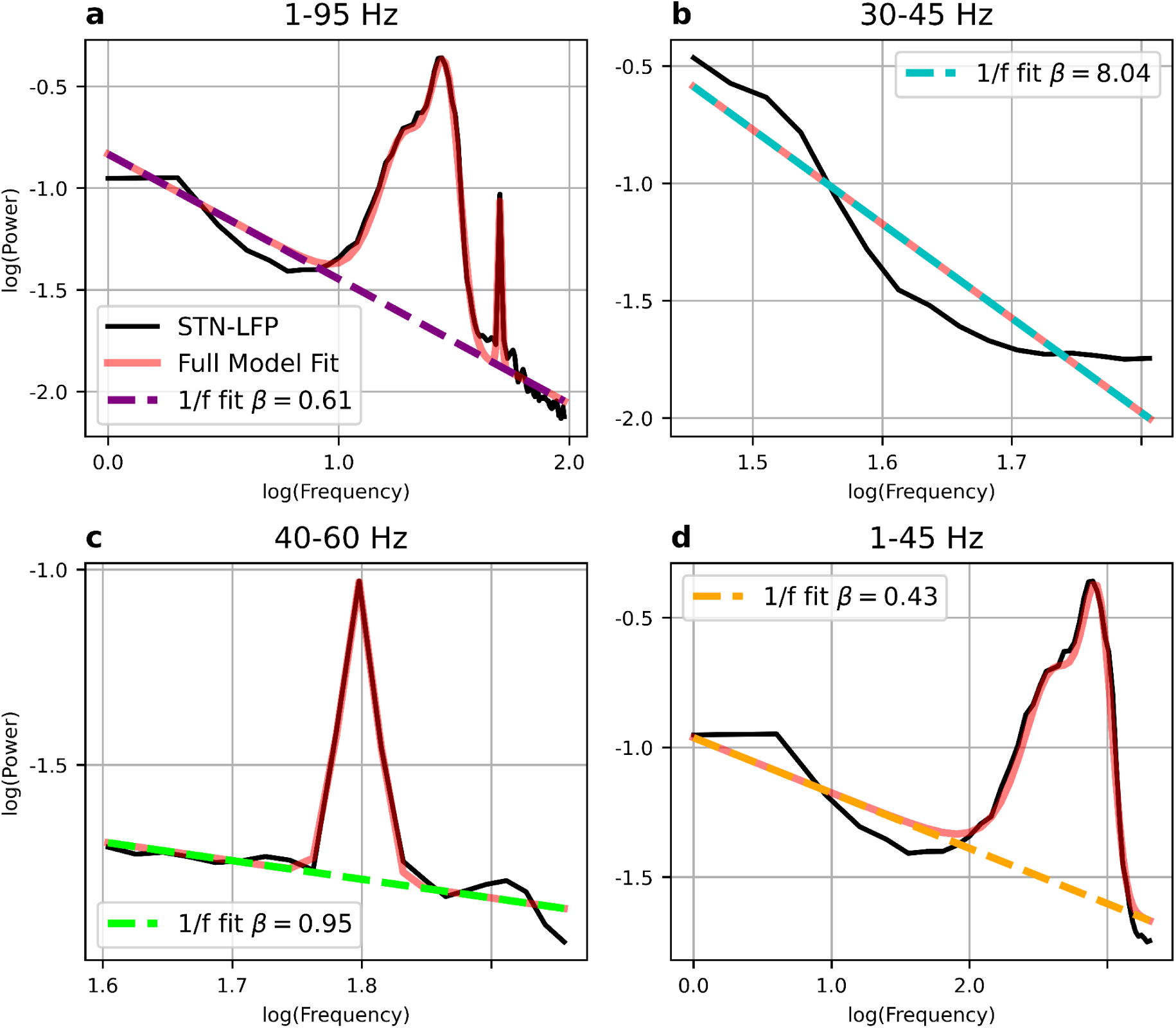
Oscillations crossing fitting range borders. FOOOF fits for the frequency ranges shown in Fig. 2 b). FOOOF parameters: a) max_n_peaks=0, 30–45 Hz, b) max_n_peaks=1, 40–60 Hz, peak_width_limits=(1, 100), 1–45 Hz, c) peak_width_limits=(1, 100), 1–95 Hz. Note that fooof fits the power line noise peak in a) and c) well. Supplementary to Fig. 3 b)

**SI Fig. 2.**
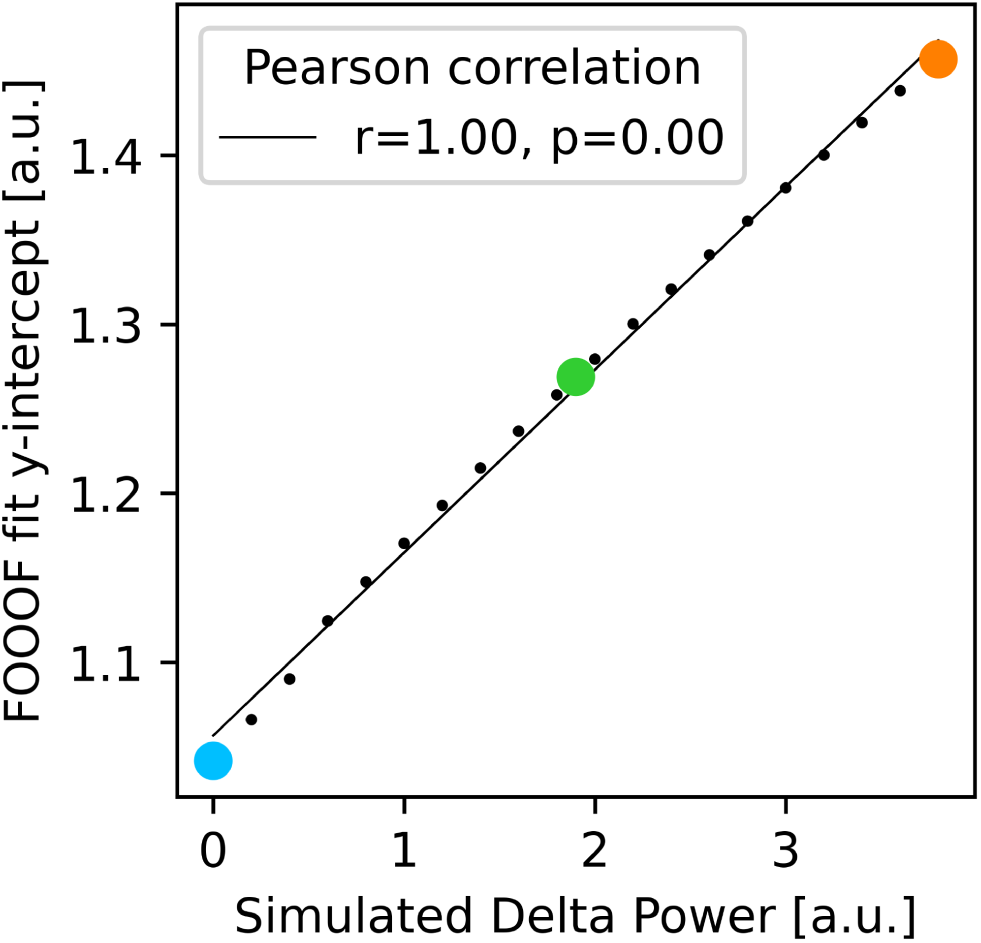
Periodic parameters impact aperiodic fits obtained with FOOOF. The y-intercepts of the FOOOF fits correlate with delta power. The blue, green, and orange data points correspond to the blue, green, and orange graphs of Fig. 3 c), the black data points indicate additional simulations.

**SI Fig. 3.**
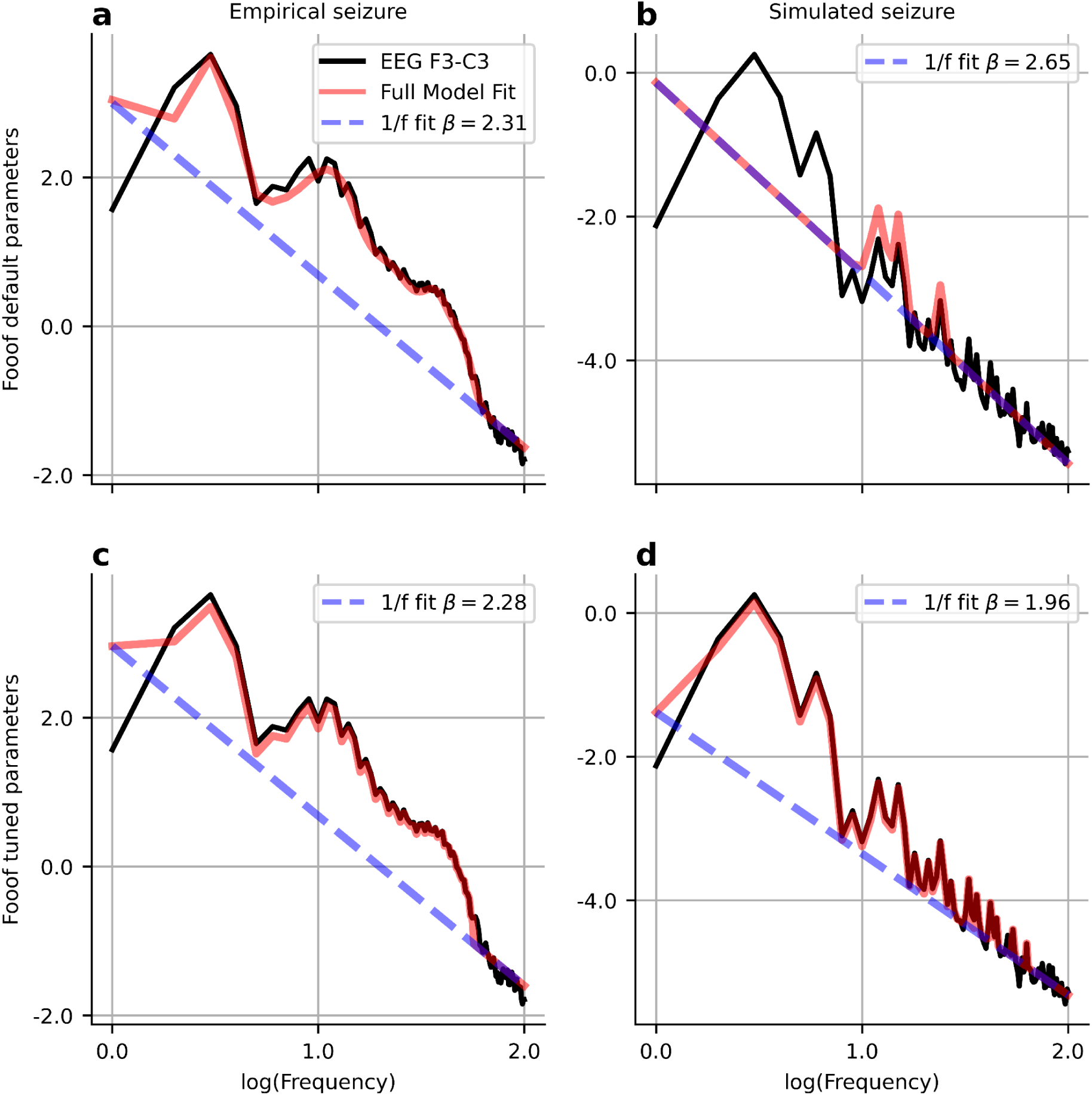
Overlapping peaks. a) FOOOF models the 3 Hz harmonics in the absence seizure recording as four peaks with center frequencies as 11 Hz, 22 Hz, 37 Hz, and 50 Hz. b) In the simulated seizure, three harmonic peaks are modeled as oscillations, whereas the rest is modeled as aperiodic component, leading to a large exponent of β =2.65 (ground truth β =1.8). c) and d) Even when the FOOOF parameters are tuned to allow maximum peak width limits of 1Hz (peak_width_limits=(0.5, 1)), FOOOF better models the harmonic peaks, but it still overestimates the 1/f exponent in the simulation. Supplementary to Fig. 4

**SI Fig. 4.**
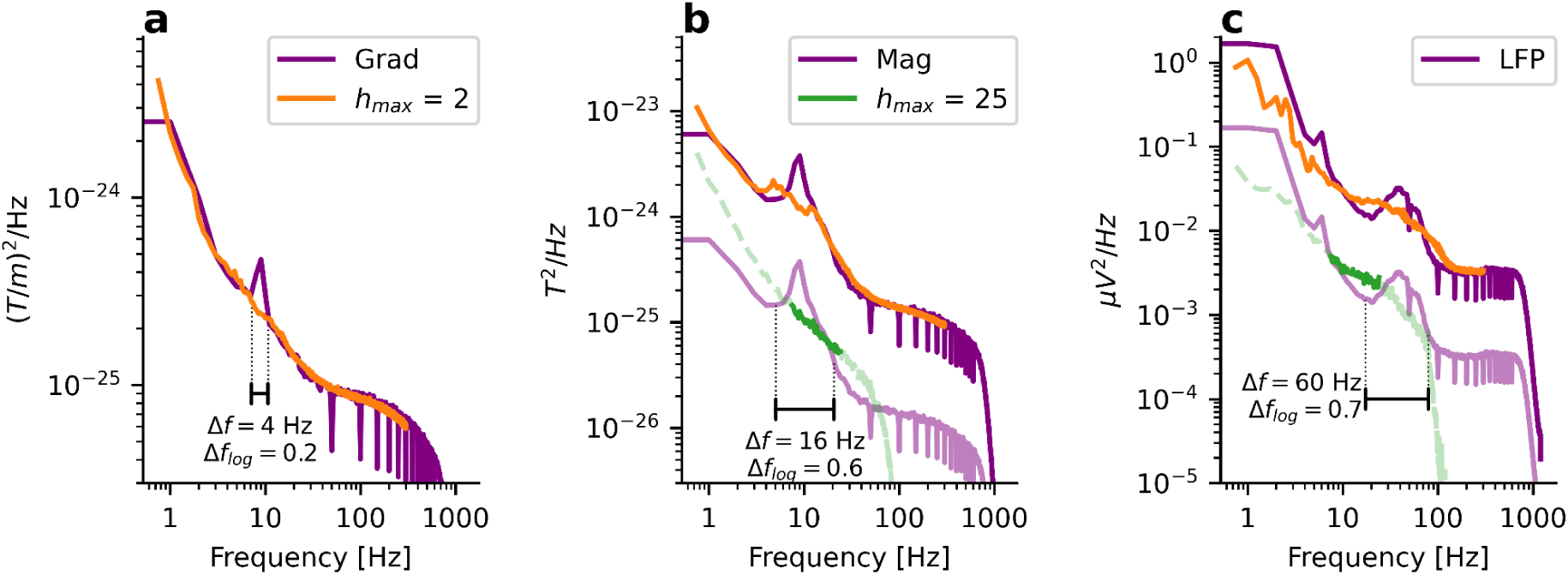
Large peak widths need large resampling factors. Dataset 3. a) MEG gradiometer PSD. A maximum resampling factor of 2 is sufficient to remove the alpha peak. b) MEG magnetometer PSD. For this peak width, a maximum resampling factor of 2 is not sufficient for removal. A maximum resampling factor of h_max_ = 25 is sufficient but leads to a large evaluated frequency range. To avoid the high-pass and spectral plateau range, only a minor part of the aperiodic component (dark green) can be used for aperiodic fitting, whereas a major part is affected by the high-pass and spectral plateau (light green dashed). c) The same holds for the large beta peak in LFP data. Note that the logarithmic peak width is essential for setting the resampling factors, not the absolute peak width. All PSDs from dataset 3. Supplementary to Fig. 6

**SI Fig. 5.**
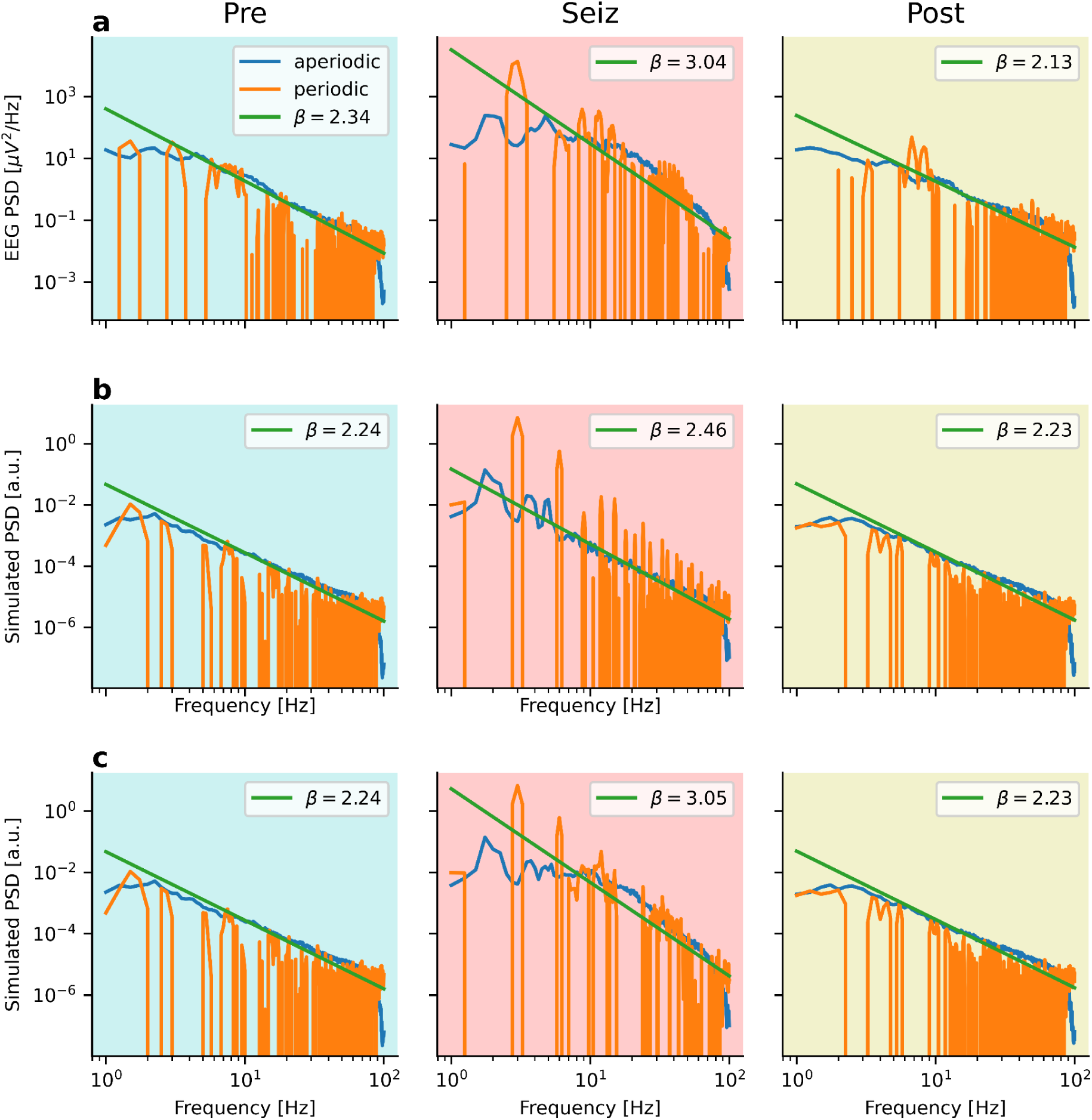
Overlapping peaks. Aperiodic (blue) and periodic (orange) extraction of IRASA and the corresponding 1/f fit (green) for the a) real and the b) and c) simulated time series in Fig. 7. For the simulation in b), IRASA can extract the harmonic 3Hz peaks well. However, the performance drops if two additional overlapping peaks are added. Supplementary to Fig. 7

## Notes

### Competing Interest Statement

The authors have declared no competing interest.

## References

Bédard, C., Kröger, H., & Destexhe, A. (2006). Does the 1/f frequency scaling of brain signals reflect self-organized critical states? Physical Review Letters, 97(11), 118102. https://doi.org/10.1103/PhysRevLett.97.118102

Belluscio, M. A., Mizuseki, K., Schmidt, R., Kempter, R., & Buzsáki, G. (2012). Cross-frequency phase-phase coupling between θ and γ oscillations in the hippocampus. The Journal of Neuroscience, 32(2), 423–435. https://doi.org/10.1523/JNEUROSCI.4122-11.2012

Bruining, H., Hardstone, R., Juarez-Martinez, E. L., Sprengers, J., Avramiea, A.-E., Simpraga, S., Houtman, S. J., Poil, S.-S., Dallares, E., Palva, S., Oranje, B., Matias Palva, J., Mansvelder, H. D., & Linkenkaer-Hansen, K. (2020). Measurement of excitation-inhibition ratio in autism spectrum disorder using critical brain dynamics. Scientific Reports, 10(1), 9195. https://doi.org/10.1038/s41598-020-65500-4

Buzsáki, G., Anastassiou, C. A., & Koch, C. (2012). The origin of extracellular fields and currents--EEG, ECoG, LFP and spikes. Nature Reviews Neuroscience, 13(6), 407–420. https://doi.org/10.1038/nrn3241

Buzsáki, G., & Draguhn, A. (2004). Neuronal oscillations in cortical networks. Science, 304(5679), 1926–1929. https://doi.org/10.1126/science.1099745

Casali, A. G., Gosseries, O., Rosanova, M., Boly, M., Sarasso, S., Casali, K. R., Casarotto, S., Bruno, M.-A., Laureys, S., Tononi, G., & Massimini, M. (2013). A theoretically based index of consciousness independent of sensory processing and behavior. Science Translational Medicine, 5(198), 198ra105. https://doi.org/10.1126/scitranslmed.3006294

Colombo, M. A., Napolitani, M., Boly, M., Gosseries, O., Casarotto, S., Rosanova, M., Brichant, J.-F., Boveroux, P., Rex, S., Laureys, S., Massimini, M., Chieregato, A., & Sarasso, S. (2019). The spectral exponent of the resting EEG indexes the presence of consciousness during unresponsiveness induced by propofol, xenon, and ketamine. NeuroImage, 189, 631–644. https://doi.org/10.1016/j.neuroimage.2019.01.024

D’Arcangelo, G., D’Antuono, M., Biagini, G., Warren, R., Tancredi, V., & Avoli, M. (2002). Thalamocortical oscillations in a genetic model of absence seizures. The European Journal of Neuroscience, 16(12), 2383–2393. https://doi.org/10.1046/j.1460-9568.2002.02411.x

Donoghue, T., Haller, M., Peterson, E. J., Varma, P., Sebastian, P., Gao, R., Noto, T., Lara, A. H., Wallis, J. D., Knight, R. T., Shestyuk, A., & Voytek, B. (2020). Parameterizing neural power spectra into periodic and aperiodic components. Nature Neuroscience, 23(12), 1655–1665. https://doi.org/10.1038/s41593-020-00744-x

Donoghue, T., Schaworonkow, N., & Voytek, B. (2021). Methodological considerations for studying neural oscillations. The European Journal of Neuroscience. https://doi.org/10.1111/ejn.15361

El Boustani, S., Marre, O., Béhuret, S., Baudot, P., Yger, P., Bal, T., Destexhe, A., & Frégnac, Y. (2009). Network-state modulation of power-law frequency-scaling in visual cortical neurons. PLoS Computational Biology, 5(9), e1000519. https://doi.org/10.1371/journal.pcbi.1000519

Engel, A. K., Fries, P., & Singer, W. (2001). Dynamic predictions: oscillations and synchrony in top-down processing. Nature Reviews Neuroscience, 2(10), 704–716. https://doi.org/10.1038/35094565

Fransson, P., Metsäranta, M., Blennow, M., Åden, U., Lagercrantz, H., & Vanhatalo, S. (2013). Early development of spatial patterns of power-law frequency scaling in FMRI resting-state and EEG data in the newborn brain. Cerebral Cortex, 23(3), 638–646. https://doi.org/10.1093/cercor/bhs047

Freeman, W. J., & Zhai, J. (2009). Simulated power spectral density (PSD) of background electrocorticogram (ECoG). Cognitive Neurodynamics, 3(1), 97–103. https://doi.org/10.1007/s11571-008-9064-y

Gao, R., Peterson, E. J., & Voytek, B. (2017). Inferring synaptic excitation/inhibition balance from field potentials. NeuroImage, 158, 70–78. https://doi.org/10.1016/j.neuroimage.2017.06.078

Gerster, M., Berner, R., Sawicki, J., Zakharova, A., Škoch, A., Hlinka, J., Lehnertz, K., & Schöll, E. (2020). FitzHugh–Nagumo oscillators on complex networks mimic epileptic-seizure-related synchronization phenomena. Chaos, 30(12), 123130. https://doi.org/10.1063/5.0021420

Gramfort, A., Luessi, M., Larson, E., Engemann, D. A., Strohmeier, D., Brodbeck, C., Goj, R., Jas, M., Brooks, T., Parkkonen, L., & Hämäläinen, M. (2013). MEG and EEG data analysis with MNE-Python. Frontiers in Neuroscience, 7, 267. https://doi.org/10.3389/fnins.2013.00267

Halgren, M., Kang, R., Voytek, B., Ulbert, I., Fabo, D., Eross, L., Wittner, L., Madsen, J., Doyle, W. K., Devinsky, O., Halgren, E., Harnett, M., & Cash, S. S. (2021). The timescale and magnitude of aperiodic activity decreases with cortical depth in humans, macaques and mice. bioRxiv. https://doi.org/10.1101/2021.07.28.454235

He, B. J. (2014). Scale-free brain activity: past, present, and future. Trends in Cognitive Sciences, 18(9), 480–487. https://doi.org/10.1016/j.tics.2014.04.003

He, B. J., Zempel, J. M., Snyder, A. Z., & Raichle, M. E. (2010). The temporal structures and functional significance of scale-free brain activity. Neuron, 66(3), 353–369. https://doi.org/10.1016/j.neuron.2010.04.020

He, W., Donoghue, T., Sowman, P. F., Seymour, R. A., Brock, J., Crain, S., Voytek, B., & Hillebrand, A. (2019). Co-Increasing Neuronal Noise and Beta Power in the Developing Brain. In bioRxiv (No. 839258). https://doi.org/10.1101/839258

Iemi, L., Busch, N. A., Laudini, A., Haegens, S., Samaha, J., Villringer, A., & Nikulin, V. V. (2019). Multiple mechanisms link prestimulus neural oscillations to sensory responses. eLife, 8. https://doi.org/10.7554/eLife.43620

Inaba, Y., D’Antuono, M., Bertazzoni, G., Biagini, G., & Avoli, M. (2009). Diminished presynaptic GABA(B) receptor function in the neocortex of a genetic model of absence epilepsy. Neuro-Signals, 17(2), 121–131. https://doi.org/10.1159/000197864

Karpova, A. V., Bikbaev, A. F., Coenen, A. M. L., & van Luijtelaar, G. (2005). Morphometric Golgi study of cortical locations in WAG/Rij rats: the cortical focus theory. Neuroscience Research, 51(2), 119–128. https://doi.org/10.1016/j.neures.2004.10.004

Kello, C. T., Brown, G. D. A., Ferrer-I-Cancho, R., Holden, J. G., Linkenkaer-Hansen, K., Rhodes, T., & Van Orden, G. C. (2010). Scaling laws in cognitive sciences. Trends in Cognitive Sciences, 14(5), 223–232. https://doi.org/10.1016/j.tics.2010.02.005

Kosciessa, J. Q., Grandy, T. H., Garrett, D. D., & Werkle-Bergner, M. (2020). Single-trial characterization of neural rhythms: Potential and challenges. NeuroImage, 206, 116331. https://doi.org/10.1016/j.neuroimage.2019.116331

Lee, D. D., & Seung, H. S. (1999). Learning the parts of objects by non-negative matrix factorization. Nature, 401(6755), 788–791. https://doi.org/10.1038/44565

Lendner, J. D., Helfrich, R. F., Mander, B. A., Romundstad, L., Lin, J. J., Walker, M. P., Larsson, P. G., & Knight, R. T. (2020). An electrophysiological marker of arousal level in humans. eLife, 9. https://doi.org/10.7554/eLife.55092

Litvak, V., Eusebio, A., Jha, A., Oostenveld, R., Barnes, G. R., Penny, W. D., Zrinzo, L., Hariz, M. I., Limousin, P., Friston, K. J., & Brown, P. (2010). Optimized beamforming for simultaneous MEG and intracranial local field potential recordings in deep brain stimulation patients. NeuroImage, 50(4), 1578–1588. https://doi.org/10.1016/j.neuroimage.2009.12.115

Litvak, V., Jha, A., Eusebio, A., Oostenveld, R., Foltynie, T., Limousin, P., Zrinzo, L., Hariz, M. I., Friston, K., & Brown, P. (2011). Resting oscillatory cortico-subthalamic connectivity in patients with Parkinson’s disease. Brain: A Journal of Neurology, 134(Pt 2), 359–374. https://doi.org/10.1093/brain/awq332

Massimini, M., Ferrarelli, F., Huber, R., Esser, S. K., Singh, H., & Tononi, G. (2005). Breakdown of cortical effective connectivity during sleep. Science, 309(5744), 2228–2232. https://doi.org/10.1126/science.1117256

Merlo, D., Mollinari, C., Inaba, Y., Cardinale, A., Rinaldi, A. M., D’Antuono, M., D’Arcangelo, G., Tancredi, V., Ragsdale, D., & Avoli, M. (2007). Reduced GABAB receptor subunit expression and paired-pulse depression in a genetic model of absence seizures. Neurobiology of Disease, 25(3), 631–641. https://doi.org/10.1016/j.nbd.2006.11.005

Miller, K. J., Sorensen, L. B., Ojemann, J. G., & den Nijs, M. (2009). Power-law scaling in the brain surface electric potential. PLoS Computational Biology, 5(12), e1000609. https://doi.org/10.1371/journal.pcbi.1000609

Miskovic, V., MacDonald, K. J., Jack Rhodes, L., & Cote, K. A. (2019). Changes in EEG multiscale entropy and power-law frequency scaling during the human sleep cycle. Human Brain Mapping, 40(2), 538–551. https://doi.org/10.1002/hbm.24393

Molina, J. L., Voytek, B., Thomas, M. L., Joshi, Y. B., Bhakta, S. G., Talledo, J. A., Swerdlow, N. R., & Light, G. A. (2020). Memantine effects on electroencephalographic measures of putative excitatory/inhibitory balance in schizophrenia. Biological Psychiatry: Cognitive Neuroscience and Neuroimaging, 5(6), 562–568. https://doi.org/10.1016/j.bpsc.2020.02.004

Muthukumaraswamy, S. D. (2013). High-frequency brain activity and muscle artifacts in MEG/EEG: a review and recommendations. Frontiers in Human Neuroscience, 7, 138. https://doi.org/10.3389/fnhum.2013.00138

Ngomba, R. T., Biagioni, F., Casciato, S., Willems-van Bree, E., Battaglia, G., Bruno, V., Nicoletti, F., & van Luijtelaar, E. L. J. M. (2005). The preferential mGlu2/3 receptor antagonist, LY341495, reduces the frequency of spike–wave discharges in the WAG/Rij rat model of absence epilepsy. Neuropharmacology, 49, 89–103. https://doi.org/10.1016/j.neuropharm.2005.05.019

Niethard, N., Hasegawa, M., Itokazu, T., Oyanedel, C. N., Born, J., & Sato, T. R. (2016). Sleep-Stage-Specific Regulation of Cortical Excitation and Inhibition. Current Biology: CB, 26(20), 2739–2749. https://doi.org/10.1016/j.cub.2016.08.035

Nikulin, V. V., Nolte, G., & Curio, G. (2011). A novel method for reliable and fast extraction of neuronal EEG/MEG oscillations on the basis of spatio-spectral decomposition. NeuroImage, 55(4), 1528–1535. https://doi.org/10.1016/j.neuroimage.2011.01.057

Onat, F. Y., van Luijtelaar, G., Nehlig, A., & Snead, O. C., 3rd. (2013). The involvement of limbic structures in typical and atypical absence epilepsy. Epilepsy Research, 103(2-3), 111–123. https://doi.org/10.1016/j.eplepsyres.2012.08.008

Peng, C. K., Havlin, S., Stanley, H. E., & Goldberger, A. L. (1995). Quantification of scaling exponents and crossover phenomena in nonstationary heartbeat time series. Chaos, 5(1), 82–87. https://doi.org/10.1063/1.166141

Pereda, E., Gamundi, A., Rial, R., & González, J. (1998). Non-linear behaviour of human EEG: fractal exponent versus correlation dimension in awake and sleep stages. Neuroscience Letters, 250(2), 91–94. https://doi.org/10.1016/s0304-3940(98)00435-2

Pumain, R., Louvel, J., Gastard, M., Kurcewicz, I., & Vergnes, M. (1992). Responses to N-methyl-D-aspartate are enhanced in rats with petit mal-like seizures. Journal of Neural Transmission. Supplementum, 35, 97–108. https://doi.org/10.1007/978-3-7091-9206-1_7

Robertson, M. M., Furlong, S., Voytek, B., Donoghue, T., Boettiger, C. A., & Sheridan, M. A. (2019). EEG power spectral slope differs by ADHD status and stimulant medication exposure in early childhood. Journal of Neurophysiology, 122(6), 2427–2437. https://doi.org/10.1152/jn.00388.2019

Romei, V., Brodbeck, V., Michel, C., Amedi, A., Pascual-Leone, A., & Thut, G. (2008). Spontaneous fluctuations in posterior alpha-band EEG activity reflect variability in excitability of human visual areas. Cerebral Cortex, 18(9), 2010–2018. https://doi.org/10.1093/cercor/bhm229

Schaworonkow, N., & Voytek, B. (2021). Longitudinal changes in aperiodic and periodic activity in electrophysiological recordings in the first seven months of life. Developmental Cognitive Neuroscience, 47, 100895. https://doi.org/10.1016/j.dcn.2020.100895

Scheer, H. J., Sander, T., & Trahms, L. (2006). The influence of amplifier, interface and biological noise on signal quality in high-resolution EEG recordings. Physiological Measurement, 27(2), 109–117. https://doi.org/10.1088/0967-3334/27/2/002

Schnitzler, A., & Gross, J. (2005). Normal and pathological oscillatory communication in the brain. Nature Reviews Neuroscience, 6(4), 285–296. https://doi.org/10.1038/nrn1650

Sharbrough, F. (1991). American Electroencephalographic Society guidelines for standard electrode position nomenclature. Journal of Clinical Neurophysiology: Official Publication of the American Electroencephalographic Society, 8(2), 200–202. https://www.ncbi.nlm.nih.gov/pubmed/2050819

Singer, W. (1999). Neuronal synchrony: a versatile code for the definition of relations? Neuron, 24(1), 49–65, 111–125. https://doi.org/10.1016/s0896-6273(00)80821-1

Stephani, T., Waterstraat, G., Haufe, S., Curio, G., Villringer, A., & Nikulin, V. V. (2020). Temporal Signatures of Criticality in Human Cortical Excitability as Probed by Early Somatosensory Responses. The Journal of Neuroscience, 40(34), 6572–6583. https://doi.org/10.1523/JNEUROSCI.0241-20.2020

Stolk, A., Brinkman, L., Vansteensel, M. J., Aarnoutse, E., Leijten, F. S., Dijkerman, C. H., Knight, R. T., de Lange, F. P., & Toni, I. (2019). Electrocorticographic dissociation of alpha and beta rhythmic activity in the human sensorimotor system. eLife, 8. https://doi.org/10.7554/eLife.48065

Tan, H. O., Reid, C. A., Single, F. N., Davies, P. J., Chiu, C., Murphy, S., Clarke, A. L., Dibbens, L., Krestel, H., Mulley, J. C., Jones, M. V., Seeburg, P. H., Sakmann, B., Berkovic, S. F., Sprengel, R., & Petrou, S. (2007). Reduced cortical inhibition in a mouse model of familial childhood absence epilepsy. Proceedings of the National Academy of Sciences of the United States of America, 104(44), 17536–17541. https://doi.org/10.1073/pnas.0708440104

Vallat, R. (2019). YASA (yet another spindle algorithm): A fast and open-source sleep spindles and slow-waves detection toolbox. Sleep Medicine, 64, S396. https://doi.org/10.1016/j.sleep.2019.11.1104

Van Veen, B. D., van Drongelen, W., Yuchtman, M., & Suzuki, A. (1997). Localization of brain electrical activity via linearly constrained minimum variance spatial filtering. IEEE Transactions on Bio-Medical Engineering, 44(9), 867–880. https://doi.org/10.1109/10.623056

Voytek, B., Kramer, M. A., Case, J., Lepage, K. Q., Tempesta, Z. R., Knight, R. T., & Gazzaley, A. (2015). Age-Related Changes in 1/f Neural Electrophysiological Noise. The Journal of Neuroscience, 35(38), 13257–13265. https://doi.org/10.1523/jneurosci.2332-14.2015

Ward, L. M. (2003). Synchronous neural oscillations and cognitive processes. Trends in Cognitive Sciences, 7(12), 553–559. https://doi.org/10.1016/j.tics.2003.10.012

Waschke, L., Donoghue, T., Fiedler, L., Smith, S., Garrett, D. D., Voytek, B., & Obleser, J. (2021). Modality-specific tracking of attention and sensory statistics in the human electrophysiological spectral exponent. In bioRxiv (p. 2021.01.13.426522). https://doi.org/10.1101/2021.01.13.426522

Waterstraat, G., Burghoff, M., Fedele, T., Nikulin, V., Scheer, H. J., & Curio, G. (2015). Non-invasive single-trial EEG detection of evoked human neocortical population spikes. NeuroImage, 105, 13–20. https://doi.org/10.1016/j.neuroimage.2014.10.024

Waterstraat, G., Fedele, T., Burghoff, M., Scheer, H.-J., & Curio, G. (2015). Recording human cortical population spikes non-invasively--An EEG tutorial. Journal of Neuroscience Methods, 250, 74–84. https://doi.org/10.1016/j.jneumeth.2014.08.013

Waterstraat, G., Körber, R., Storm, J.-H., & Curio, G. (2021). Noninvasive neuromagnetic single-trial analysis of human neocortical population spikes. Proceedings of the National Academy of Sciences of the United States of America, 118(11). https://doi.org/10.1073/pnas.2017401118

Wen, H., & Liu, Z. (2016). Separating Fractal and Oscillatory Components in the Power Spectrum of Neurophysiological Signal. Brain Topography, 29(1), 13–26. https://doi.org/10.1007/s10548-015-0448-0

Zanos, T. P., Mineault, P. J., & Pack, C. C. (2011). Removal of spurious correlations between spikes and local field potentials. Journal of Neurophysiology, 105(1), 474–486. https://doi.org/10.1152/jn.00642.2010

